# Single-molecule imaging reveals the roles of the membrane-binding motif and the C-terminal domain of RNase E in its localization and diffusion in *Escherichia coli*

**DOI:** 10.1101/2024.11.05.622141

**Authors:** Laura Troyer, Yu-Huan Wang, Shobhna Shobhna, Seunghyeon Kim, Brooke Ramsey, Jeechul Woo, Emad Tajkhorshid, Sangjin Kim

## Abstract

In *Escherichia coli*, RNase E, a central enzyme in RNA processing and mRNA degradation, contains a catalytic N-terminal domain (NTD), a membrane-targeting sequence (MTS), and a C-terminal domain (CTD). We investigated how MTS and CTD influence RNase E localization, diffusion, and function. Super-resolution microscopy revealed that ∼93% of RNase E localizes to the inner membrane and exhibits slow diffusion similar to polysomes. Comparing the native amphipathic MTS with a transmembrane motif showed that the MTS confers slower diffusion and stronger membrane binding. The CTD further slows diffusion by increasing mass but unexpectedly weakens membrane association. RNase E mutants with partial cytoplasmic localization displayed enhanced co-transcriptional degradation of *lacZ* mRNA. These findings indicate that variations in the MTS and the presence of the CTD shape the spatiotemporal organization of RNA processing in bacterial cells, providing mechanistic insight into how RNase E domain architecture influences its cellular function.

## INTRODUCTION

RNase E (RNE) is the main endoribonuclease in *Escherichia coli*, known for its role in RNA processing and mRNA degradation^1–3^. It is an essential protein^4,5^, and homologous proteins are found across many bacterial species^6–8^. The essentiality stems from the N-terminal domain (NTD), or the catalytic domain^9^. The NTD is followed by a membrane targeting sequence (MTS) and the C-terminal domain (CTD), or macromolecular interaction domain^2^, where RhlB (a DEAD-box RNA helicase), PNPase (a 3’→5’ exonuclease), and enolase (a glycolytic enzyme) bind to form the RNA degradosome complex^6^. The MTS forms an amphipathic α helix, responsible for the localization of RNE on the inner membrane^10,11^. Interestingly, the membrane localization of RNE and the presence of the CTD are not essential in *E. coli* nor are they conserved across bacterial species, in contrast to the broad conservation of the NTD across bacteria as well as chloroplasts^7,12,13^. This raises a question about the roles of membrane localization and the CTD in the *in vivo* function of RNE.

*E. coli* strains with cytoplasmic RNE (due to the removal of the MTS) are viable, although they grow more slowly than the wild-type (WT) cells^10,14^. *In vitro* studies have shown that membrane binding of RNE does not necessarily increase its enzymatic activity^10,14^. However, membrane localization is likely important for gene regulation *in vivo* because RNE becomes sequestered from the cytoplasmic pool of mRNAs, giving mRNAs time for translation. This idea is supported by our recent observation that the membrane-bound RNE limits the degradation of nascent mRNAs while cytoplasmic RNE (ΔMTS) can degrade nascent mRNAs during transcription^15^. We found that transcripts encoding membrane proteins can be an exception to this rule, in that they can experience co-transcriptional degradation assisted by the transertion effect^15^. These findings agree with results from a genome-wide study, indicating that the membrane localization of RNE allows for differential regulation of mRNA stability for genes encoding cytoplasmic proteins versus inner membrane proteins in *E. coli*^16^.

Previous studies have reported evidence that *E. coli* RNE can localize in the cytoplasm—for example, when cells were grown anaerobically^17^ or when membrane fluidity was reduced by changes in lipid composition^18^. These findings imply that RNE can dissociate from the membrane; however, the origin of its weak membrane binding remains unknown.

Across bacteria, several species within α-proteobacteria have cytoplasmic RNE^19^ while other species possess membrane-bound RNE. Among these, *B. subtilis* RNase Y (a functional homolog of RNE) associates with the membrane via a transmembrane (TM) motif^20^, instead of an amphipathic motif used by *E. coli* and other γ-proteobacteria^7^. Given the diversity of membrane-binding motifs that have arisen through evolution, it should be possible to engineer *E. coli* RNE with a TM motif. Such a mutant would provide a useful model for investigating the impact of membrane-binding motifs on the localization, diffusion, and activity of RNE.

Lastly, the CTD of *E. coli* RNE is an intrinsically disordered region^21^ that, while nonessential for cell viability, enhances the enzymatic activity of the NTD^15,22–24^. As the primary binding site for degradosome components^6^, the CTD is thought to facilitate mRNA degradation by recruiting these proteins near the catalytic NTD. However, our recent study showed that these associated proteins have minimal impact on the degradation rate of *lacZ* mRNA, whereas deletion of the CTD markedly stabilizes the transcript^15^, suggesting a possible intramolecular allosteric effect within RNE. In the present study, we further show that the CTD modulates the membrane binding affinity of RNE, possibly by affecting its conformation. These results reveal a previously underappreciated role of the CTD in regulating RNE function beyond degradosome assembly, with implications for how the spatial organization and structural dynamics of RNE fine-tune RNA degradation in bacteria.

In this study, we quantified the membrane binding percentage (MB%) of RNE in *E. coli* using single-molecule microscopy and showed that membrane association governs its diffusion and mRNA-degradation activity. Perturbing the native MTS, substituting it with LacY TM segments, and deleting the CTD collectively revealed how the MTS and CTD set RNE’s spatial organization and provided routes for tuning activity through subcellular control.

## RESULTS

### Membrane binding percentage (MB%) of RNE

First, we investigated the subcellular localization of RNE in live cells. Previous fluorescence microscopy studies have shown that RNE is localized to the inner membrane in *E. coli*^10,11,16^, but the percentage of membrane-bound molecules has not been quantitatively examined in live cells. To address this gap, we fused RNE with a photo-convertible fluorescent protein, mEos3.2^25^ and imaged individual RNE molecules over time in two dimensions (**Fig. 1A**). The positions of fluorescent molecules were identified in each frame and linked into trajectories using the open-source software u-track^26^ (**Fig. 1A**). In this section, we analyze localization, and diffusion dynamics are addressed in subsequent sections.

**Figure 1:**
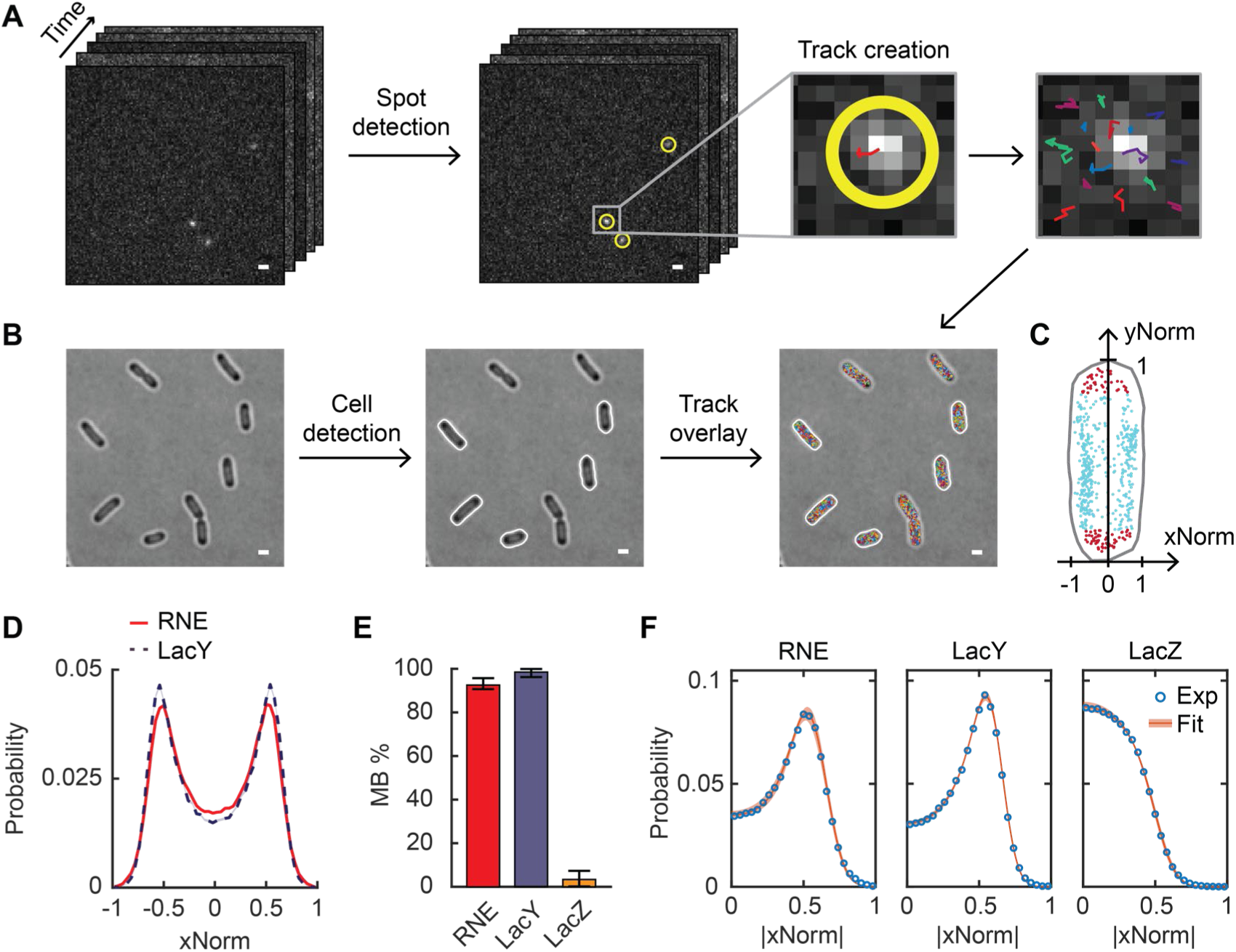
Analysis of single-molecule images for the subcellular localization and dynamics of proteins. (**A**) Single-molecule image analysis. Spots were detected in each frame (highlighted with yellow circles), and tracks were created across frames (different colors were chosen for different tracks). (**B**) Cell detection. Cell outlines were determined from bright-field images. Only non-dividing cells were analyzed (indicated by white outlines). (**C**) Normalized position of spots of RNE along the short (x) and the long (y) axes of an example cell. Red spots are inside the cell endcaps, and cyan spots are in the cylindrical region of the cell. (**D**) xNorm histogram of RNE and LacY. Only spots in the cylindrical region of cells (like cyan spots in **C**) were included, totaling n = 143,000 spots. The standard error of the mean (SEM) calculated from bootstrapping is displayed as a shaded area but is smaller than the line width (see **Fig. S1A** for details). (**E**) The membrane binding percentage (MB%) of RNE, LacY, and LacZ. Error bars are from the 95% confidence interval. (**F**) Histogram of absolute xNorm and model fitting of RNE, LacY, and LacZ to determine MB%. Orange highlights indicate the range of xNorm expected based on the standard deviations in the parameter values estimated by MCMC. The white scale bars in panels **A-B** are 1 µm. See **Table S6** for data statistics.

Subcellular locations were calculated relative to the cell boundaries identified from bright-field images using another open-source image analysis package Oufti^27^ (**Fig 1B**). To combine data from many cells, molecular positions along the short and long axes of a cell were normalized to the cell width and cell length, yielding xNorm and yNorm, respectively (**Fig. 1C**). Based on yNorm, molecules within the cylindrical part of the cell were selected, and their xNorm values were used to generate an xNorm histogram. Hereinafter, we focus on the xNorm histogram to compare the membrane enrichment across protein constructs.

The xNorm histogram of RNE shows two peaks corresponding to the inner membrane on each side of the cell (**Fig. 1D**, **Fig. S1A**), very similar to the xNorm histogram of LacY, obtained by imaging LacY-mEos3.2 using the same method. LacY is a membrane channel for lactose and is composed of 12 TM segments^28^. It is expected to be inserted into the inner membrane during translation^29–31^, such that all imaged LacY is expected to be localized in the inner membrane^32^.

We further quantified the percentage of molecules bound to the membrane (membrane-binding percentage or MB%) from the xNorm histogram. For this analysis, we first confirmed that proteins localized on the membrane and in the cytoplasm are detected with equal probability, despite differences in their mobilities (**Fig. S1B-C**). Next, we developed a mathematical model based on a 2D projection of molecules randomly distributed either on the surface of or within a cylinder. The model includes imaging effects that affect the shape of xNorm histograms: localization error, the limited focal depth of the quasi-TIRF illumination we used, and the location of the inner membrane relative to the cell boundary (**Fig. S1D-G**). Model fitting was performed using a Markov-Chain Monte Carlo algorithm (MCMC). For validation, we applied the model to xNorm histograms of LacY and LacZ, which serve as benchmarks for complete membrane binding and complete cytoplasmic localization, respectively. The MB% of LacY was 99% with a 95% confidence interval of [96%, 100%], and LacZ showed MB% of 3.4% [0.2%, 7.3%] (**Fig. 1E-F**). Both MB% agree with the expectations for membrane and cytoplasmic proteins. For RNE, we found an MB% of 93% [91%, 96%] (**Fig. 1E**). The xNorm histogram of the fastest 7% of the RNE population (based on the diffusion coefficient, as discussed below) exhibited a cytoplasmic localization pattern (without the two membrane-associated peaks), supporting the existence of a cytoplasmic RNE subpopulation (**Fig. S2A**).

We confirmed that the MTS (residue 568-582) is essential for the membrane binding of RNE, as deletion of the MTS sequence made the xNorm like that of LacZ (**Fig. 2A-B**, **Fig. S2D**). We note that the xNorm profile of RNE ΔMTS was slightly different from that of LacZ near the center line of the cell (x = 0), suggesting fewer RNE ΔMTS molecules were at the midline of the cell. Also, xNorm fitting yielded an MB% of 33%. These results may reflect nucleoid exclusion of RNE ΔMTS due to its large size^33^. Consistently, a smaller cytoplasmic RNE variant generated by CTD truncation (RNE (1-529) or RNE ΔMTS ΔCTD) exhibited a xNorm profile closer to that of LacZ (**Fig. 2B**, **Fig. S2E**).

**Figure 2:**
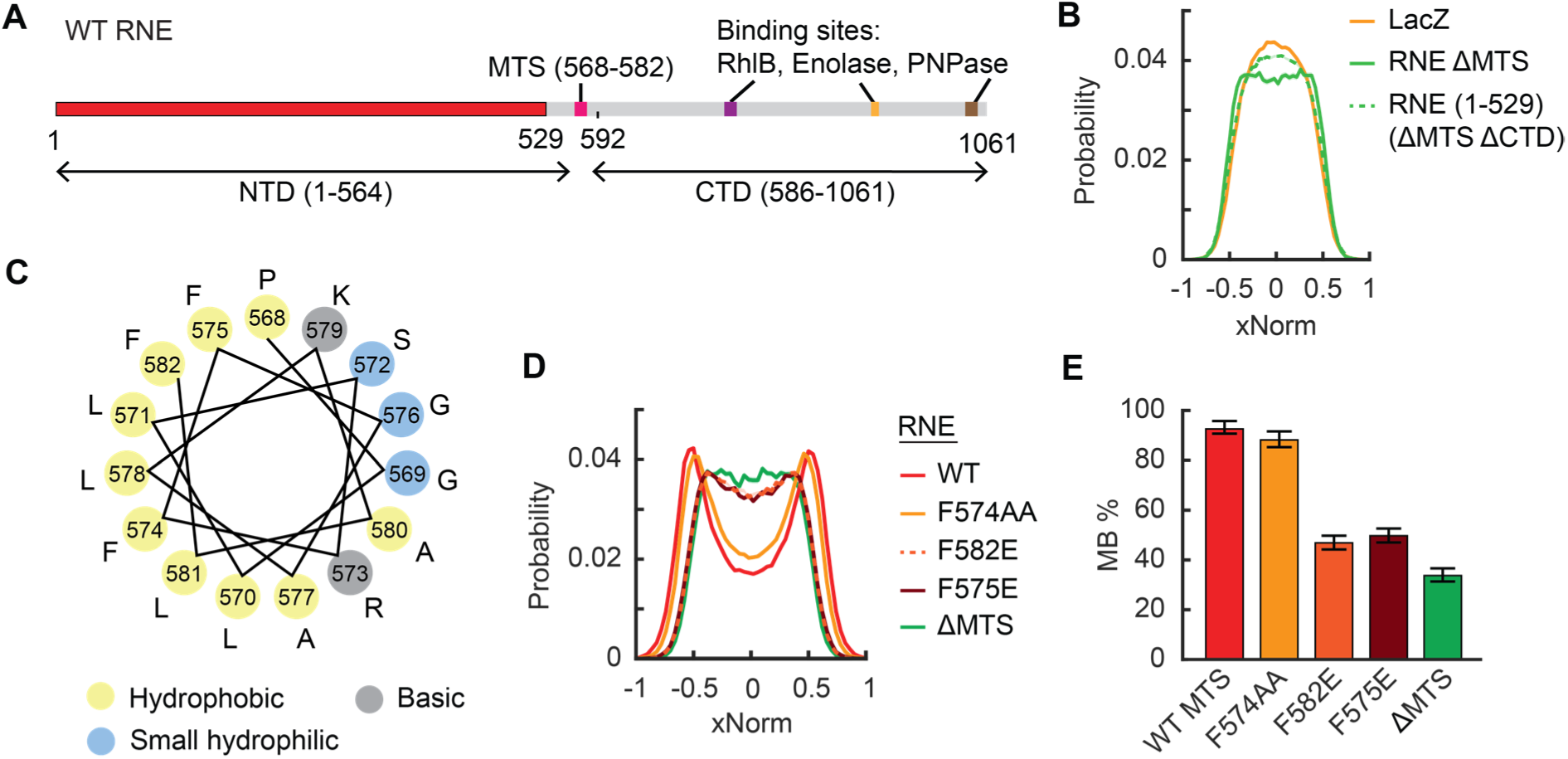
Mutations in the MTS affecting the localization of RNE. (**A**) Linear representation of the RNE monomer. The NTD and the CTD are defined as regions flanking the MTS. Numbers indicate the amino acid residues. (**B**) xNorm histograms of cytoplasmic RNE mutants. (**C**) Helical wheel diagram of the MTS region of RNE (residue 568-582). (**D**) xNorm histogram of RNE MTS point mutants. (**E**) MB% of RNE MTS point mutants. Error bars indicate the 95% confidence interval. In panels B and D, the SEM from bootstrapping is shown but is smaller than the line width. See **Table S6** for data statistics.

Is there a critical residue(s) within the MTS required for membrane binding? The MTS forms an amphipathic α-helix, in which hydrophobic residues are expected to align on one side of the helix (**Fig. 2C**)^10^. Previous studies suggested that replacing one of the hydrophobic residues with a hydrophilic amino acid can disrupt membrane binding of RNE^10,11^. We revisited the point mutations discussed in Khemici et al^10^ and analyzed MB% by mEos3.2 imaging. We found that the F574A F575A double mutation (noted as F574AA) did not significantly affect MB%, consistent with the fact that the substituted amino acids remain hydrophobic (**Fig. 2D-E**, **Fig. S2F**). In contrast, F582E and F575E mutations reduced MB% to 47% and 50%, respectively (**Fig. 2E**), and their xNorm histograms resembled that of RNE ΔMTS, suggesting predominantly cytoplasmic localization (**Fig. 2D**, **Fig. S2D**, **S2G-H**). These findings indicate that the phenylalanine residues at positions 575 and 582 are critical for membrane association of RNE.

### Effect of membrane binding on the diffusion of RNE

The membrane localization of RNE likely limits its diffusion and interaction with mRNA targets. Additionally, interaction with ribosome-bound mRNAs can further slow the diffusion of RNE. Here, we examined how these factors contribute to the diffusion dynamics of RNE.

To measure the diffusion of RNE, we analyzed the trajectories of individual RNE-mEos3.2 imaged at a 21.7-ms acquisition interval (**Fig. 1A**). We calculated the diffusion coefficient *D* by fitting the mean-squared displacement (MSD) of each trajectory to the equation MSD = 4*D*τ + *b*, where τ is lag time and *b* accounts for both dynamic and static localization errors^34^ (**Fig. 3A**). We obtained *D*_RNE_ = 0.0184 ± 0.0002 µm^2^/s (mean ± SEM). This value was well above the lower detection limit of our microscope, determined using stationary, surface-immobilized mEos3.2, *D* = 0.0020 ± 0.0001 μm^2^/s (**Fig. S3A, Supplementary Discussion**). Notably, *D*_RNE_ was comparable to that of ribosome-bound mRNAs, estimated to be *D* ∼0.015 µm^2^/s based on the diffusion of ribosomal protein L1 (**Fig. S3B**).

**Figure 3:**
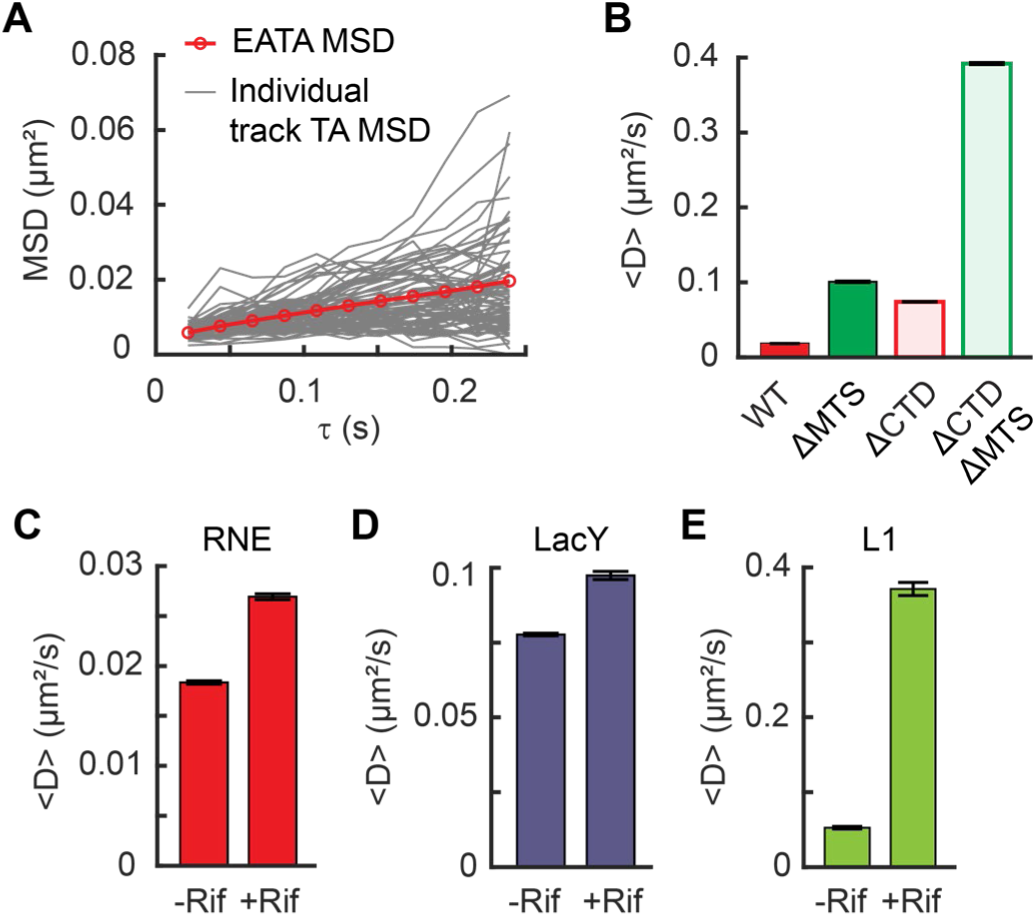
RNE diffusion, influenced by membrane binding and interactions with mRNAs. (**A**) MSD versus time delay (τ) for RNE. Ensemble-averaged time-averaged (EATA) MSD was calculated by averaging the time-averaged MSD of individual tracks. (**B**) Mean diffusion coefficients of various RNE mutants, lacking the MTS and/or the CTD. (**C-E**) Change in the mean diffusion coefficient of RNE (**C**), LacY (**D**), and ribosome L1 protein (**E**) when cellular RNAs were depleted by rifampicin treatment. Error bars in panels B-E represent the SEM. See **Table S6** for data statistics.

To assess how membrane association affects diffusion, we compared *D* of WT RNE and the ΔMTS mutant. These two proteins have similar molecular masses, as the MTS comprises only 15 of the 1061 residues in a monomer of RNE (**Fig. 2A**). Therefore, any difference in *D* can be attributed to their subcellular localizations (membrane vs cytoplasm) rather than mass. We found that *D*_ΔMTS_ is ∼5.5 times that of *D*_RNE_ (**Fig. 3B**).

We examined another pair of RNE mutants that differ in localization (membrane vs cytoplasm) but are similar in size: membrane-bound RNE (1-592) and cytoplasmic RNE (1-529), both lacking the CTD. These truncated variants diffused faster than their full-length counterparts due to reduced mass, but their *D* values still differed by a factor of ∼5.3 due to localization (**Fig. 3B**). Together, these results suggest that the membrane binding reduces RNE mobility by a factor of 5.

### Diffusion of RNE in the absence of mRNA substrates

When some of RNE molecules interact with mRNA, their diffusion can slow due to the added mass of mRNA and ribosomes, possibly yielding slower RNE subpopulations. Such mobility-based subpopulations have been observed for RNA polymerases and ribosomes in *E. coli* and used to estimate the fraction of molecules interacting with RNA^35–37^.

To test the effect of mRNA substrates on RNE diffusion, we treated cells with rifampicin (rif), which blocks transcription initiation, thus depleting cellular mRNAs^38^. In rif-treated cells, *D*_RNE_ increased to 0.0270 ± 0.0003 µm^2^/s, which is 1.47 ± 0.010 times that in untreated cells (**Fig. 3C**). We note that *D*_LacY_ also increased to 1.25 ± 0.01 times relative to untreated cells (**Fig. 3D**). Although this increase is relatively small, the increase in *D*_LacY_ was unexpected because LacY is not an RNA-binding protein. The increase in *D*_LacY_ likely results from the depletion of mRNAs near the membrane (e.g., the mRNAs undergoing transertion), which could otherwise hinder the diffusion of membrane proteins^39,40^. The similar fold change in *D*_RNE_ and *D*_LacY_ upon rif treatment suggests that the change in RNE diffusion may largely be attributed to physical changes in the intracellular environment (such as reduced viscosity or macromolecular crowding^41,42^), rather than a loss of RNA-RNE interactions.

Because the rif-induced change in *D*_RNE_ is largely physical, we next examined why eliminating RNA-RNE interactions does not further increase RNE mobility, using the ribosome as a benchmark. The diffusion of ribosomal protein L1 became 7 times as fast as that in rif-treated cells (**Fig. 3E**), consistent with previous reports^35,36^. This big change can be explained by the fact that ribosomes form polysomes, where the effective mass of L1 protein in untreated cells would be 2 or more times that in rif-treated cells (where it remains as a free subunit). In the case of RNE, it forms the RNA degradosome complex, whose mass can be from 450 kDa^6^ to 2.3 MDa depending on the occupancy of the RNA degradosome component proteins (RhlB, PNPase, enolase)^2,43–46^ (**Supplementary Discussion)**. Even at its largest size, the RNE complex is smaller than a 70S ribosome (∼2.5 MDa^47^). This means that if RNE interacts with an mRNA associated with n ribosomes, the total mass of the RNE complex would increase by a factor of n or more. Thus, a substantial increase in *D*_RNE_ would be expected upon mRNA depletion, assuming that a significant fraction of RNE is engaged with mRNA-ribosome assemblies.

To explain the marginal increase in *D*_RNE_ upon rif treatment, we considered two possibilities; (1) only a small percentage of RNE molecules interacts with mRNAs at a given time and/or (2) RNE interacts with mRNAs only briefly, unlike ribosomes which spend an order of 10-100 s in a polysome state during translation elongation^48^. To distinguish between these possibilities, we attempted to increase the cellular pool of polysomes, either by treating cells with a translation elongation inhibitor chloramphenicol^37^ or by overexpressing *lacZ* mRNA from a high-copy plasmid (**Fig. S3D-E**). In both cases, a larger fraction of RNE would engage with mRNAs, potentially increasing the fraction of RNE in the slow-diffusing state. However, *D*_RNE_ remained unchanged compared to untreated cells (**Fig. S3F**). This result rules out the possibility that only a small percentage of RNE interacts with mRNAs and instead weighs in favor of the scenario that RNE-mRNA interactions are brief. Specifically, if RNE interacts with mRNAs for ∼20 ms or less, the slow-diffusing state would last shorter than the frame interval and remain undetected in our experiment.

### Diffusion and localization of the MTS and TM segments

Unlike RNE in *E. coli*, RNase Y, a functional homolog of RNE in *B. subtilis*, is localized to the membrane via a TM domain^20^. We wondered if there are differences between a peripheral motif (like the MTS of *E. coli*’s RNE) and a TM motif in terms of membrane localization and mobility. To address this question, we created RNE mutants in which the MTS was replaced with a TM domain. For the TM domain, we used TM segments of LacY, a native *E. coli* protein, rather than using the TM motif from *B. subtilis* RNase Y, which might interact with the *E. coli* membrane in a non-native manner.

Before creating the RNE mutants, we characterized the MB% and the diffusion of individual short membrane-binding motifs. Native LacY contains 12 TM segments, arranged into two groups of six^28^. We successfully expressed constructs containing the first two TM segments (LacY (1-74) or LacY2), the first six TM segments (LacY (1-193) or LacY6), and the full-length LacY (LacY12), each fused to mEos3.2 and expressed from a chromosomal IPTG-inducible promoter (**Fig. 4A**). We then imaged their membrane localization and diffusion. Both the MTS segment and LacY-derived TM segments showed a strong membrane enrichment (**Fig. 4B**). In terms of diffusion, LacY2 and LacY6 diffused faster than the MTS segment (**Fig. 4C**), contrary to expectations based on size (or mass)-dependent diffusion (**Fig. 4D**).

**Figure 4:**
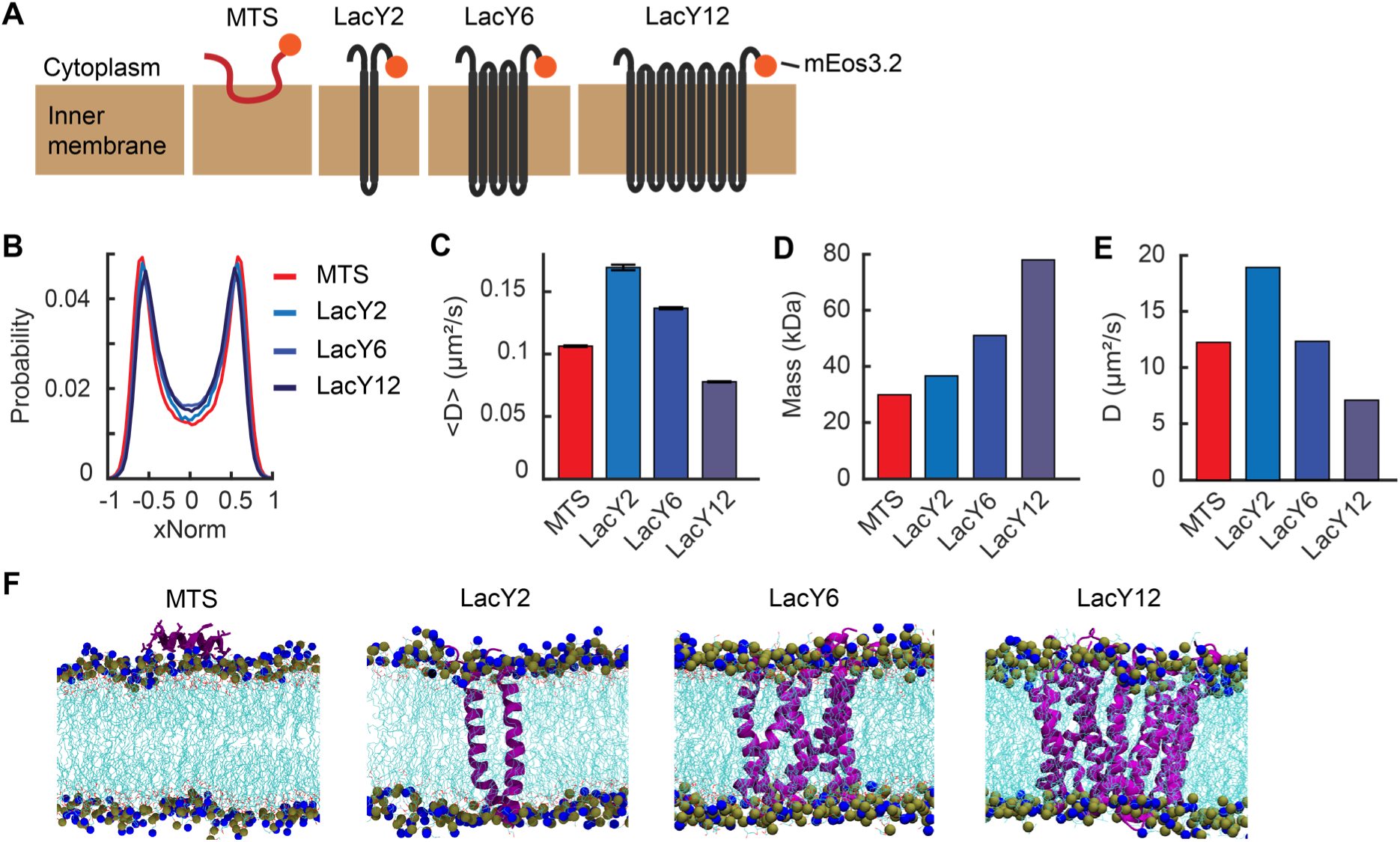
Localization and diffusion of membrane-binding motifs. (**A**) Cartoon schematic of the membrane-binding motifs used in this study (not to scale). The orange circles indicate mEos3.2 used for imaging. (**B**) xNorm histograms of membrane-binding motifs. The SEM from bootstrapping is displayed but smaller than the line width. Data are from at least 107,000 spots. (**C**) Mean diffusion coefficients of membrane-binding motifs. Error bars are the SEM from at least 3,000 tracks. (**D**) Estimated mass of membrane-binding motifs based on the amino acid sequence including linkers and mEos3.2. (**E**) Diffusion coefficients of the membrane-binding motifs obtained from all-atom MD simulation. (**F**) Representative simulation snapshots of the membrane-binding motifs embedded in the *E. coli* membrane. Proteins are displayed in purple, and lipid tails are shown in cyan. Nitrogen and phosphorus atoms of the lipid head groups are represented in the van der Waals form in blue and grey, respectively. See **Table S6** for data statistics.

According to the Stokes-Einstein relation for diffusion in simple fluids^49^ and the Saffman-Delbruck diffusion model for membrane proteins^50^, *D* decreases as particle size increases, albeit with different scaling behaviors. Specifically, the Saffman-Delbruck model predicts that *D* for membrane proteins decreases logarithmically with increasing radius of the membrane-embedded region, assuming a constant membrane environment^50^. Thus, if size (or mass) were the primary determinant of diffusion, LacY2 and LacY6 would diffuse more slowly than the smaller MTS. The observed discrepancy instead implies that *D* may be governed by how each motif interacts with the membrane. For example, the way that TM domains are anchored to the membrane may facilitate faster lateral diffusion with surrounding lipids.

Despite the prevalence of peripheral membrane proteins^51^, how they interact with the membrane and how this differs from TM proteins remain poorly understood. To further explore this, we conducted all-atom molecular dynamics (MD) simulations of the MTS and the LacY variants interacting with the *E. coli* membrane using the NAMD software^52^. In the simulations, protein motion was calculated for 1 µs. Although the absolute *D* values were higher than experimental values (possibly due to the absence of mEos3.2 in the model), the overall trends were preserved; among the LacY series, larger constructs diffused more slowly (**Fig. 4E-F**, **Fig. S4A**). Most importantly, the MTS again diffused more slowly than LacY2 *in silico* (**Fig. 4E**, **Fig. S4A**). By calculating membrane-protein interaction energies, we found that the MTS-membrane interactions were more stable than those of LacY2 (**Fig. S4B**). These results suggest that the slower diffusion of the MTS is due to stronger interactions with lipid head groups compared to membrane-embedded TM segments.

### RNE mutants carrying a TM motif

Since LacY2 and LacY6 showed strong membrane enrichment similar to LacY12 (**Fig. 4B**), we replaced the MTS in RNE with LacY2, LacY6, and LacY12 in the presence or absence of the CTD (**Fig. 5A-B** and **Fig. S5**). All chimeric RNE mutants were expressed from the native chromosomal locus as the only copy of *rne*, with mEos3.2 fused at the C terminus for imaging. The resulting strains exhibited no noticeable differences in growth rate compared to the WT strain (**Table S3**), suggesting that the RNE mutants were functionally active.

**Figure 5:**
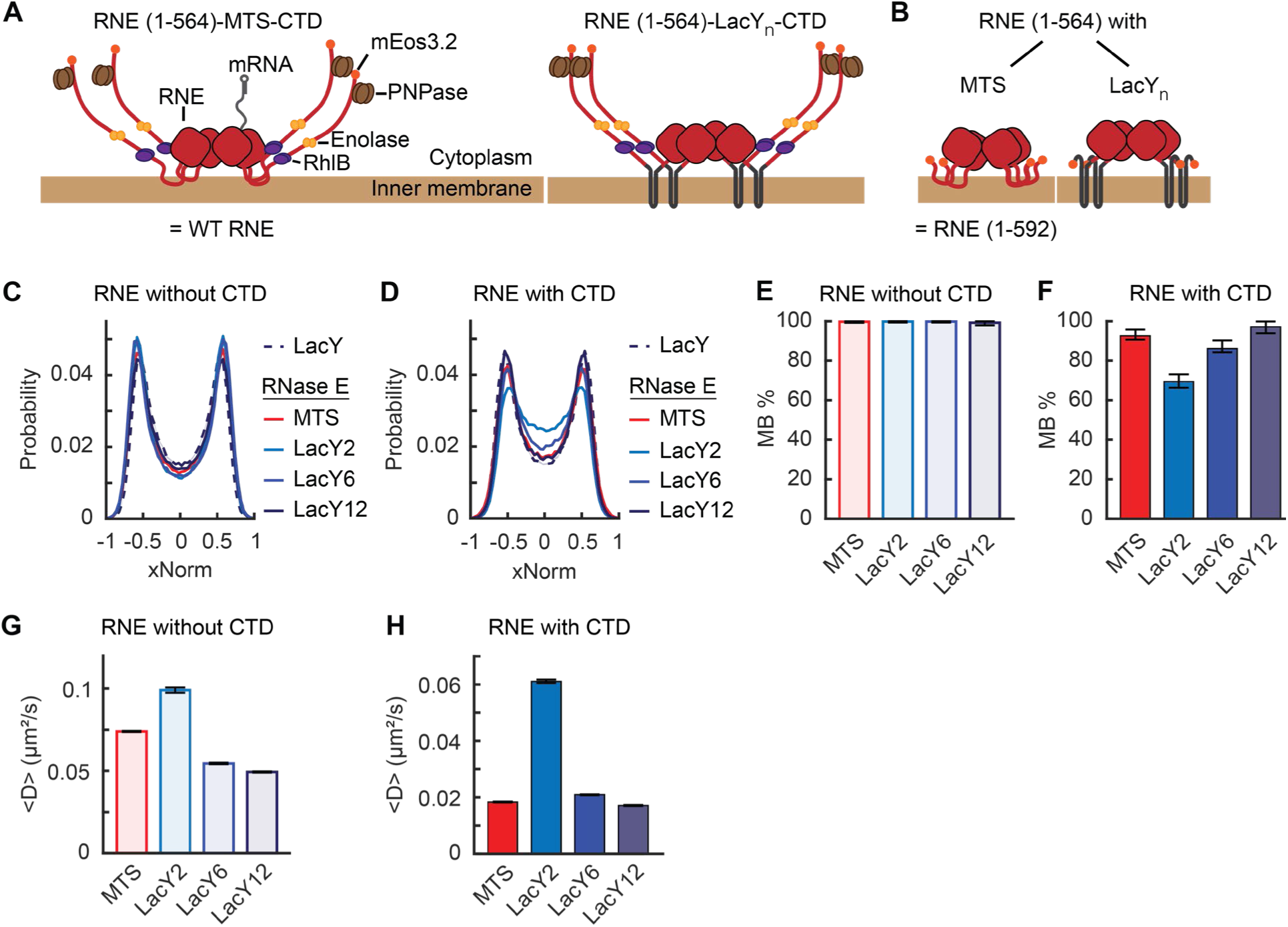
Localization and diffusion of chimeric RNE with or without the CTD. (**A-B**) Cartoon schematic of RNE chimeric variants with the CTD (**A**) and without the CTD (**B**). They are not to scale. (**C-D**) xNorm histograms of chimeric RNE localization compared with that of LacY. The SEM from bootstrapping is displayed but smaller than the line width. (**E-F**) MB% of chimeric RNE mutants without the CTD (**E**) or with the CTD (**F**) with various membrane-binding motifs. Error bars are from a 95% confidence interval. (**G-H**) Mean diffusion coefficients of chimeric RNE without the CTD (**G**) or with the CTD (**H**). Error bars are the SEM. Each data set contains at least 70,000 tracks for diffusion or 72,000 spots for xNorm. See **Table S6** for data statistics.

xNorm histograms of the ΔCTD mutants indicated membrane localization similar to LacY (**Fig. 5C**). However, mutants containing the CTD showed noticeable cytoplasmic subpopulations when LacY2 and LacY6 were used in place of the MTS (**Fig. 5D**, **Fig. S2J-K**, **Fig. S6B**). Mathematical model fitting of the xNorm histograms estimated the MB% of RNE-LacY2-CTD and RNE-LacY6-CTD to be 69% [66%, 73%] and 86% [84%, 90%], respectively (**Fig. 5F**). We note that imperfect membrane localization was observed only in mutants containing the CTD; the same protein without the CTD showed MB% of 100% (**Fig. 5E-F, Fig. S6A-B**). These findings suggest that the CTD may contribute to unstable membrane binding of RNE. Supporting this idea, previously characterized RNE MTS point mutants with MB% <50% (**Fig. 2D-E**) also exhibited increased MB% upon the CTD removal (**Fig. S6A-B**). Such a difference in MB% between the CTD-containing and the CTD-lacking mutants was not observed in the chimera based on LacY12 (**Fig. 5E-F**), possibly due to the stable membrane insertion by LacY12.

Next, we examined whether the *D* of chimeric RNE mutants varied depending on the type of membrane-binding motifs. For example, the MTS diffused more slowly than LacY2 and LacY6, despite being smaller in size (**Fig. 4C-D**). Based on this, we expected the chimeric RNE with LacY2 or LacY6 to diffuse faster than RNE with MTS. Indeed, in the absence of the CTD, we found that the *D* of LacY2-based RNE was 1.33 ± 0.01 times as fast as the MTS-based RNE (**Fig. 5G**). However, LacY6-based RNE did not diffuse faster than the MTS-based version (**Fig. 5G**). This result may be due to the high TM load (24 helices) created by four LacY6 anchors in the RNE tetramer. Although all constructs are tetrameric, the 24-helix load (LacY6), compared with 8 (LacY2) and 4 (MTS), likely enlarges the membrane-embedded footprint and increases drag, thereby changing the mobility advantages assessed as standalone membrane anchors.

In the presence of the CTD, the *D* of LacY2 and LacY6-based RNE became 3.33 ± 0.004 and 1.14 ± 0.01 times that of the MTS-based RNE counterpart (i.e. the WT RNE), respectively (**Fig. 5H**). This is likely influenced by the presence of a cytoplasmic population (∼31% for LacY2 and ∼14% for LacY6; **Fig. 5F**, **Fig S2J-K**), which diffuses more rapidly than membrane-bound molecules (possibly by a factor of five, based on **Fig. 3B**). Taken together, our data suggest that the CTD weakens the membrane association of RNE and small TM motifs can facilitate the diffusion of RNE.

### Functional consequence of subcellular localization and diffusion of RNE

To check the functional consequence of cytoplasmic localization of RNE, we measured *lacZ* mRNA degradation in various RNE mutants presented in this study. Recently, we developed an assay to quantify both co-transcriptional and post-transcriptional degradation rates of *lacZ* mRNA by inducing its transcription for only 75 s, thereby capturing the degradation of nascent mRNA^15^. In WT cells, we found that the co-transcriptional degradation rate (*k*_d1_) is about 10 times slower than the post-transcriptional degradation rate (*k*_d2_)^15^. In cells expressing RNE ΔMTS, however, *k*_d1_ increases by a factor of ∼3, suggesting that cytoplasmic RNE can freely diffuse and degrade nascent mRNAs^15^. This result led us to hypothesize that RNE variants exhibiting a cytoplasmic subpopulation (**Fig. 2E** and **5F**) may exhibit a larger *k*_d1_ compared to more membrane-bound variants.

We repeated this assay in a strain expressing the WT RNE fused to mEos3.2 (**Fig. S7A**). Note that this strain is different from the one we used for imaging because a monocistronic *lacZ* gene is needed for the transient induction assay^15^. The relative abundances of 5’ *lacZ* mRNA (Z5) remained constant prior to the rise in 3’ *lacZ* mRNA (Z3) levels (between ∼100 and 210 s), confirming negligible co-transcriptional degradation in WT cells^15^ (**Fig. 6A**). However, in cells expressing RNE-LacY2-CTD (MB% = 69%), Z5 levels exhibited a downward trend during the same time window (**Fig. 6B**). The estimated *k*_d1_ was close to what was observed in RNE ΔMTS^15^ (gray line, *p* = 0.28; **Fig. 6C**). A similarly high *k*_d1_ was also observed in RNE MTS point mutations, F582E and F757E, which exhibited a ΔMTS-like xNorm profile (**Fig. 2D**), supporting the idea that the cytoplasmic subpopulation of RNE enables co-transcriptional mRNA degradation in *E. coli* (**Fig. 6C**). Notably, RNE-LacY6-CTD, which also exhibited a cytoplasmic subpopulation (∼14% from MB% of 86%), did not exhibit a significant increase in *k*_d1_ (**Fig. 6C**), suggesting a critical amount of cytoplasmic population may be needed to facilitate co-transcriptional mRNA degradation

**Figure 6:**
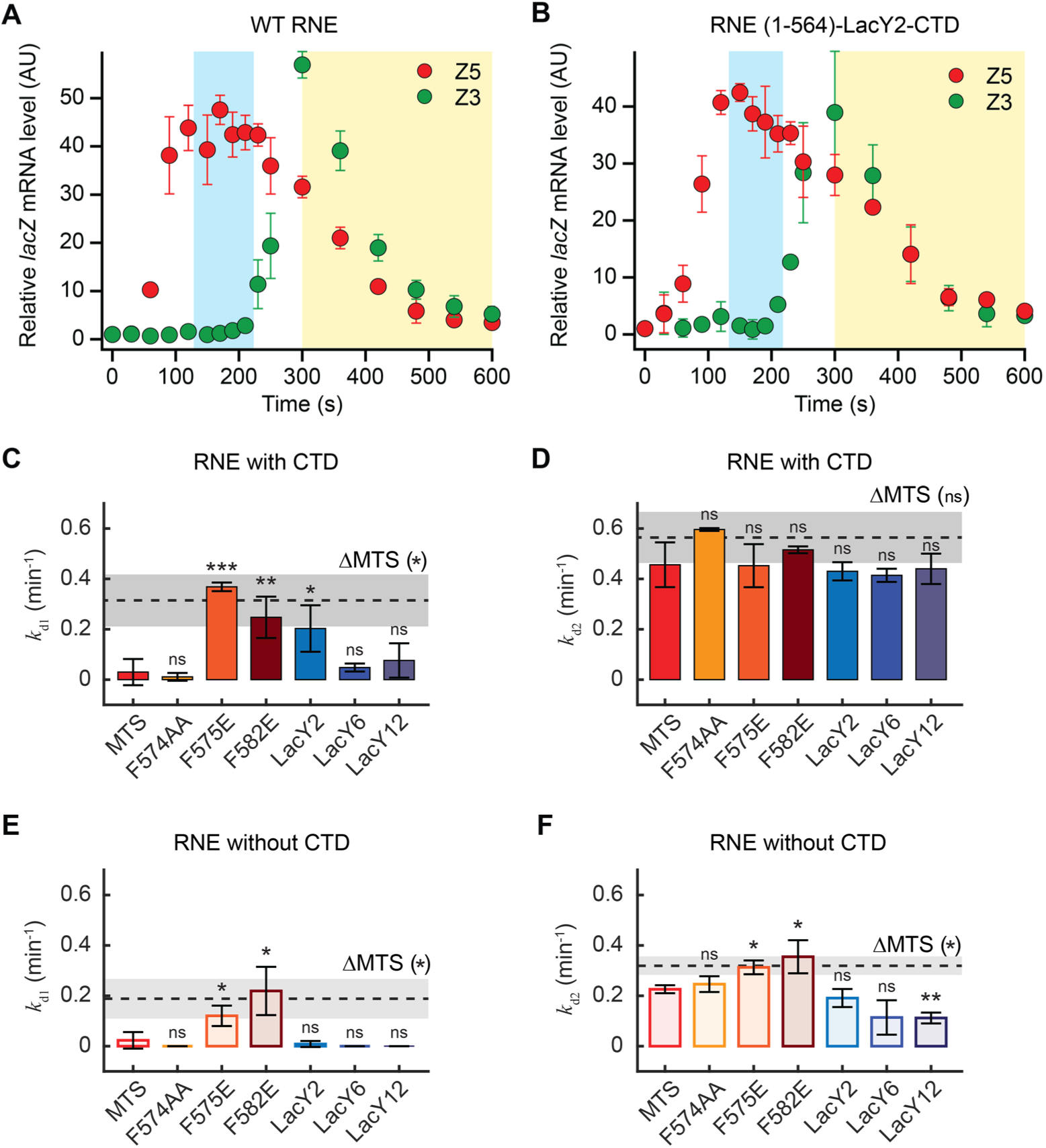
*lacZ* mRNA degradation rates in RNE mutant strains. (**A-B**) *lacZ* mRNA levels in WT RNE (**A**, strain SK595) and in RNE-LacY2-CTD (**B**, strain SK505) when *lacZ* transcription was induced with 0.2 mM IPTG at t = 0 s and re-repressed with 500 mM glucose at t = 75 s. Blue and yellow regions indicate the time windows used to measure k_d1_ and k_d2_, respectively, by exponential fitting of 5’ *lacZ* mRNA (Z5) in individual replicates. (**C-F**) Co-transcriptional and post-transcriptional *lacZ* mRNA degradation rates, k_d1_ (**C, E**) and k_d2_ (**D, F**), respectively, in various RNE mutants containing different membrane-binding motifs, either with the CTD (solid bars, **C-D**) or ΔCTD (light bars, **E-F**). The dotted lines indicate the k_d1_ and k_d2_ values of cytoplasmic RNE ΔMTS (strain SK339 in **C-D**)_15_ or RNE ΔMTS ΔCTD (strain SK370, in **E-F**). In all panels, error bars represent the standard deviations from 2-3 biological replicates. Two-sample *t-*tests were performed relative to the MTS case in each graph (See **Table S7** for the *p* values).

We next examined whether the post-transcriptional mRNA degradation rate (*k*_d2_) is limited by the slow diffusion of membrane-bound RNE. In the presence of the CTD, *k*_d2_ did not significantly vary across membrane-targeting variants (**Fig. 6D**). For example, even though the LacY12 motif slows RNE diffusion, its *k*_d 2_ was similar to that of WT RNE (**Fig. 6D**). A critical control for this comparison is RNE abundance: because RNE autoregulates its expression by degrading its own transcript^53^, slower diffusion could elevate RNE levels and mask the negative effect of the reduced mobility. We tested this explicitly and found that over-expression of WT RNE does not significantly change *k*_d1_ or *k*_d2_ (**Fig. S7B**). Thus, when the CTD is present, neither copy number nor diffusion of membrane-bound RNE limits *k*_d2_.

In the absence of the CTD, *k*_d1_ was high in RNE variant F582E (MB% = 67%; **Fig. S6A**), reaching levels close to its cytoplasmic counterpart, the ΔMTS ΔCTD mutant (gray line, *p* = 0.7; **Fig. 6E**). *k*_d2_ values for ΔCTD variants were, in general, lower than those of their CTD-containing counterparts (**Fig. 6F**), indicating the importance of the CTD for the catalytic activity. These findings are consistent with previous reports on RNE constructs based on the MTS^15,22–24^. Among ΔCTD variants with different membrane motifs, point mutants F575E and F582E (MB% = 91% and 67%, respectively; **Fig. S6A**) exhibited higher *k*_d2_ than the MTS-based variant, in agreement with the fact that completely cytoplasmic ΔMTS ΔCTD exhibited higher *k*_d2_ than the MTS-containing ΔCTD (**Fig. 6F**). However, the LacY2-based RNE variant, which diffuses faster than the MTS version (**Fig. 5G**), did not show a corresponding increase in *k*_d2_ (**Fig. 6F**). Plus, LacY6 and LacY12 versions showed even lower *k*_d2_ than MTS-based RNE ΔCTD. Overall, these results indicate that the reduced *k*_d2_ caused by ΔCTD cannot be rescued by faster diffusion of membrane-bound RNE. In fact, it may be further impaired by large and slow membrane motifs (such as LacY12). The presence of the CTD appears to buffer the effects of large membrane-binding motifs on RNE’s catalytic activity, helping to maintain efficient post-transcriptional mRNA degradation (*k*_d2_).

## DISCUSSION

Our study establishes the membrane enrichment and slow diffusion of RNE in *E. coli*. This supports the notion that sequestration of RNE on the membrane confers the spatial and temporal separation between synthesis and decay of mRNAs^15^. Furthermore, the processing of rRNA^54–58^ and tRNA^59,60^ and small RNA-based gene regulations^61,62^ mediated by RNE likely take place on the membrane.

For WT RNE, our analysis showed MB% of 93%, close to 91% previously reported using immunogold labeling and freeze-fracture electron microscopy^63^. Whether the MB% of RNE changes under different growth conditions remains to be tested. As a case study, we examined the MB% of WT RNE when cells were grown in M9 minimal medium with succinate as the sole carbon source without supplements, a condition different from that used in our primary experiments (M9 glycerol with supplements). The MTS segment exhibited a reduction in MB% from 100% to 83% (**Fig. S6C**), suggesting that MTS-mediated membrane binding can be sensitive to growth conditions. However, the MB% of WT RNE was only marginally affected by the same media change (**Fig. S6C**). We speculate that tetramer formation stabilizes RNE membrane localization, even when individual MTS motifs exhibit a weak membrane affinity. A similar phenomenon has been observed with MinD; although its amphipathic motif alone is cytoplasmic, MinD becomes membrane-associated upon dimerization, indicating that oligomerization enhances the membrane binding of this peripheral membrane protein^64^. By analogy, the tetrameric structure of RNE may reinforce membrane association, buffering against changes in cellular metabolism or lipid environment that would otherwise weaken MTS-mediated binding.

While individual TM motifs exhibited strong membrane association regardless of growth conditions (**Fig. S6D**), we found LacY2-based RNE can lose membrane binding affinity, unlike MTS-based RNE. The loss of MB% in LacY2-based RNE was observed only in the presence of the CTD (**Fig. S6D**), suggesting that the CTD negatively affects membrane binding of RNE, possibly by altering protein conformation. In fact, all ΔCTD RNE mutants we tested exhibited higher MB% than their CTD-containing counterparts (**Fig. S6A-B**). For example, WT RNE (containing CTD) showed an MB% of 93%, whereas its ΔCTD version, RNE (1-592), showed 100%. Similarly, the chimeric RNE-LacY6-CTD showed an MB% of 86%, while its ΔCTD version showed 100% (**Fig. 5E-F**, **Fig. S6A-B**). A similar trend was observed for MTS point mutants (**Fig. S6A-B**), further supporting that the CTD decreases membrane association across RNE variants. We speculate that this effect may be related to the CTD’s role in promoting phase-separated ribonucleoprotein condensates, as observed in *Caulobacter crescentus*^19^. In *E. coli*, we also observed a modest increase in the clustering tendency of RNE compared to ΔCTD (**Fig. S8**).

The *D*_RNE_ we measured in *E. coli* (0.018 μm^2^/s) is comparable to those measured in other bacterial species. For example, in *C. crescentus*, the *D* of its cytoplasmic RNE was shown to be about 0.03 μm^2^/s^65^. The diffusion of RNase Y in *B. subtilis* was found in either a slow (0.031 μm^2^/s) or fast (0.3 μm^2^/s) population^66^, with the slow population corresponding to RNase Y bound to mRNA and/or the putative RNA degradosome and the fast population representing freely diffusing RNase Y. In the case of *E. coli* RNE, a two-population fit of the *D* histogram based on MB% identified the slow and fast subpopulations whose *D* values differed by a factor of 4 (**Fig. S3C**). This agrees with our finding that cytoplasmic RNE ΔMTS diffuses ∼5 times as fast as WT RNE on average (**Fig. 3B**).

Diffusion is affected by the size of the particle in a given medium^50^, and it has been used to identify different size forms of a protein due to biochemical interactions or complex formation^36,67^. Related, RNA substrates interacting with RNE can also increase the effective mass of RNE and lower its *D* value. However, when cellular mRNAs were depleted by rifampicin treatment, RNE diffusion increased less than that of other RNA-binding proteins related to transcription and translation. For example, the large and small ribosomal subunits^35,36,68^, tRNA^69^, and RNA polymerase^37^ showed a large (about 10-20 fold) increase in *D* upon rifampicin treatment. Interestingly, Hfq, an RNA chaperon involved in small RNA regulation together with RNE, showed a moderate increase (∼2 fold) in *D* when RNA was depleted by rifampicin^70^. We note that our result is consistent with previous studies that examined the effect of rifampicin on RNE diffusion in *E. coli*^11,71^ as well as RNE in *C. crescentus*^65^ and RNase Y in *B. subtilis*^66,72^. One possible explanation is that RNA-bound RNE (and RNase Y) is short-lived compared to our frame interval (∼20 ms), unlike other RNA-binding proteins related to transcription and translation, interacting with RNA for ∼1 min for elongation^48^.

Lastly, the slow diffusion of the MTS in comparison to LacY2 and LacY6 suggests that MTS is less favorable for rapid lateral motion in the membrane. Our MD simulations demonstrated that this reduced diffusivity may originate from a stronger interaction energy between the protein and the membrane. Whether this property is specific to the MTS or generalizable to other peripheral versus integral membrane motifs remains to be tested. We speculate that this can be a general phenomenon, as peripheral motifs interact with lipids orthogonally while integral membrane motifs align parallel to the bilayer and may diffuse more freely by coupling with the motion of surrounding lipids. The diffusion behavior of membrane-bound proteins reflects underlying protein-lipid interactions and membrane dynamics^73^. Accordingly, future work may analyze how protein-lipid interaction strength and membrane dynamics contribute to differences in lateral diffusion between peripheral and integral membrane proteins.

Altogether, our work highlights strategies to modulate the MB% and diffusion of RNE to possibly affect mRNA degradation rates. For example, membrane-bound RNE lacking the CTD can stabilize mRNAs and increase protein expression. This idea has been used in a commercial *E. coli* BL21 strain (Invitrogen’s One Shot BL21 Star) to increase recombinant protein yield^22^. Additionally, RNE variants with environmentally responsive MB%, such as RNE-LacY2-CTD, could be harnessed to tune mRNA half-lives and protein expression levels under different growth conditions.

## ACKNOWLEDGMENTS

We thank Drs. Mark Arbing, Agamemnon Carpousis, Johan Elf, and Christine Jacobs-Wagner for strains. We thank Maggie Liu, Kavya Vaidya, and Zach Wang for their contributions in the early phase of this work and the members of Kim lab for critical reading of the manuscript. This work was supported by the NSF Center for Physics of Living Cells (1430124), NSF Science and Technology Center for Quantitative Cell Biology (2243257), NIH (R35GM143203; R24GM145965), and Searle Scholars Program.

## AUTHOR CONTRIBUTIONS

L.T. and S.K. designed the research; L.T. and Y.W. performed imaging and analysis; L.T. and Se.K. measured mRNA lifetimes; LT, Se.K., and S.K. performed genetics; L.T., Y.W., and J.W. developed data analysis methods; S. performed all-atom MD simulations under supervision of E.T.; B.R. performed epi-fluorescence imaging; L.T. and S.K. wrote the manuscript with input from all authors; S.K. supervised the study.

## REFERENCES

1 Mudd, E. A., Krisch, H. M. & Higgins, C. F. RNase E, an endoribonuclease, has a general role in the chemical decay of Escherichia coli mRNA: evidence that rne and ams are the same genetic locus. Molecular Microbiology 4, 2127–2135 (1990). 10.1111/j.1365-2958.1990.tb00574.x

2 Mackie, G. A. RNase E: at the interface of bacterial RNA processing and decay. Nature Reviews Microbiology 11, 45–57 (2013). 10.1038/nrmicro2930

3 Hui, M. P., Foley, P. L. & Belasco, J. G. Messenger RNA degradation in bacterial cells. Annual Review of Genetics 48, 537–559 (2014). 10.1146/annurev-genet-120213-092340

4 Apirion, D. & Lassar, A. B. A conditional lethal mutant of Escherichia coli which affects the processing of ribosomal RNA. Journal of Biological Chemistry 253, 1738–1742 (1978). 10.1016/S0021-9258(17)34927-X

5 Babitzke, P. & Kushner, S. R. The Ams (altered mRNA stability) protein and ribonuclease E are encoded by the same structural gene of Escherichia coli. Proceedings of the National Academy of Sciences 88, 1–5 (1991). doi:10.1073/pnas.88.1.1

6 Carpousis, A. J. The RNA degradosome of Escherichia coli: an mRNA-degrading machine assembled on RNase E. Annual Review of Microbiology 61, 71–87 (2007). 10.1146/annurev.micro.61.080706.093440

7 Aït-Bara, S. & Carpousis, A. J. RNA degradosomes in bacteria and chloroplasts: classification, distribution and evolution of RNase E homologs. Molecular Microbiology 97, 1021–1135 (2015). 10.1111/mmi.13095

8 Mardle, C. E. et al. A structural and biochemical comparison of ribonuclease E homologues from pathogenic bacteria highlights species-specific properties. Scientific Reports 9, 7952 (2019). 10.1038/s41598-019-44385-y

9 Callaghan, A. J. et al. Structure of Escherichia coli RNase E catalytic domain and implications for RNA turnover. Nature 437, 1187–1191 (2005). 10.1038/nature04084

10 Khemici, V., Poljak, L., Luisi, B. F. & Carpousis, A. J. The RNase E of Escherichia coli is a membrane-binding protein. Molecular Microbiology 70, 799–813 (2008). 10.1111/j.1365-2958.2008.06454.x

11 Strahl, H. et al. Membrane recognition and dynamics of the RNA degradosome. PLOS Genetics 11, e1004961 (2015). 10.1371/journal.pgen.1004961

12 Kaberdin, V. R. et al. The endoribonucleolytic N-terminal half of Escherichia coli RNase E is evolutionarily conserved in Synechocystis sp. and other bacteria but not the C-terminal half, which is sufficient for degradosome assembly. Proceedings of the National Academy of Sciences 95, 11637–11642 (1998). doi:10.1073/pnas.95.20.11637

13 Lee, K. & Cohen, S. N. A Streptomyces coelicolor functional orthologue of Escherichia coli RNase E shows shuffling of catalytic and PNPase-binding domains. Molecular Microbiology 48, 349–360 (2003). 10.1046/j.1365-2958.2003.03435.x

14 Hadjeras, L. et al. Detachment of the RNA degradosome from the inner membrane of Escherichia coli results in a global slowdown of mRNA degradation, proteolysis of RNase E and increased turnover of ribosome-free transcripts. Molecular Microbiology 111, 1715–1731 (2019). 10.1111/mmi.14248

15 Kim, S., Wang, Y.-H., Hassan, A. & Kim, S. Re-defining how mRNA degradation is coordinated with transcription and translation in bacteria. bioRxiv (2024). 10.1101/2024.04.18.588412

16 Moffitt, J. R., Pandey, S., Boettiger, A. N., Wang, S. & Zhuang, X. Spatial organization shapes the turnover of a bacterial transcriptome. eLife 5, e13065 (2016). 10.7554/eLife.13065

17 Murashko, O. N. & Lin-Chao, S. Escherichia coli responds to environmental changes using enolasic degradosomes and stabilized DicF sRNA to alter cellular morphology. Proceedings of the National Academy of Sciences 114, E8025–E8034 (2017). doi:10.1073/pnas.1703731114

18 Gohrbandt, M. et al. Low membrane fluidity triggers lipid phase separation and protein segregation in living bacteria. The EMBO Journal 41, e109800 (2022). 10.15252/embj.2021109800

19 Al-Husini, N., Tomares, D. T., Bitar, O., Childers, W. S. & Schrader, J. M. α-Proteobacterial RNA degradosomes assemble liquid-liquid phase-separated RNP bodies. Molecular Cell 71, 1027–1039.e1014 (2018). 10.1016/j.molcel.2018.08.003

20 Lehnik-Habrink, M. et al. RNase Y in Bacillus subtilis: a natively disordered protein that is the functional equivalent of RNase E from Escherichia coli. Journal of Bacteriology 193, 5431–5441 (2011). doi:10.1128/jb.05500-11

21 Callaghan, A. J. et al. Studies of the RNA degradosome-organizing domain of the Escherichia coli ribonuclease RNase E. Journal of Molecular Biology 340, 965–979 (2004). 10.1016/j.jmb.2004.05.046

22 Lopez, P. J., Marchand, I., Joyce, S. A. & Dreyfus, M. The C-terminal half of RNase E, which organizes the Escherichia coli degradosome, participates in mRNA degradation but not rRNA processing in vivo. Molecular Microbiology 33, 188–199 (1999). 10.1046/j.1365-2958.1999.01465.x

23 Leroy, A., Vanzo, N. F., Sousa, S., Dreyfus, M. & Carpousis, A. J. Function in Escherichia coli of the non-catalytic part of RNase E: role in the degradation of ribosome-free mRNA. Molecular Microbiology 45, 1231–1243 (2002). 10.1046/j.1365-2958.2002.03104.x

24 Islam, M. S., Bandyra, K. J., Chao, Y., Vogel, J. & Luisi, B. F. Impact of pseudouridylation, substrate fold, and degradosome organization on the endonuclease activity of RNase E. RNA 27, 1339–1352 (2021). 10.1261/rna.078840.121

25 Zhang, M. et al. Rational design of true monomeric and bright photoactivatable fluorescent proteins. Nature Methods 9, 727–729 (2012). 10.1038/nmeth.2021

26 Jaqaman, K. et al. Robust single-particle tracking in live-cell time-lapse sequences. Nature Methods 5, 695–702 (2008). 10.1038/nmeth.1237

27 Paintdakhi, A. et al. Oufti: an integrated software package for high-accuracy, high-throughput quantitative microscopy analysis. Molecular Microbiology 99, 767–777 (2016). 10.1111/mmi.13264

28 Abramson, J. et al. Structure and mechanism of the lactose permease of Escherichia coli. Science 301, 610–615 (2003). doi:10.1126/science.1088196

29 Ahrem, B., Hoffschulte, H. K. & Müller, M. In vitro membrane assembly of a polytopic, transmembrane protein results in an enzymatically active conformation. J Cell Biol 108, 1637–1646 (1989). 10.1083/jcb.108.5.1637

30 Nagamori, S., Vázquez-Ibar, J. L., Weinglass, A. B. & Kaback, H. R. In vitro synthesis of lactose permease to probe the mechanism of membrane insertion and folding. Journal of Biological Chemistry 278, 14820–14826 (2003). 10.1074/jbc.M300332200

31 Stochaj, U. & Ehring, R. The N-terminal region of Escherichia coli lactose permease mediates membrane contact of the nascent polypeptide chain. European Journal of Biochemistry 163, 653–658 (1987). 10.1111/j.1432-1033.1987.tb10914.x

32 Volkov, I. L. et al. Spatiotemporal kinetics of the SRP pathway in live E. coli cells. Proceedings of the National Academy of Sciences 119, e2204038119 (2022). doi:10.1073/pnas.2204038119

33 Thappeta, Y. et al. Glycogen phase separation drives macromolecular rearrangement and asymmetric division in E. coli. bioRxiv (2024). 10.1101/2024.04.19.590186

34 Savin, T. & Doyle, P. S. Static and dynamic errors in particle tracking microrheology. Biophysical Journal 88, 623–638 (2005). 10.1529/biophysj.104.042457

35 Bakshi, S., Siryaporn, A., Goulian, M. & Weisshaar, J. C. Superresolution imaging of ribosomes and RNA polymerase in live Escherichia coli cells. Molecular Microbiology 85, 21–38 (2012). 10.1111/j.1365-2958.2012.08081.x

36 Sanamrad, A. et al. Single-particle tracking reveals that free ribosomal subunits are not excluded from the Escherichia coli nucleoid. Proceedings of the National Academy of Sciences 111, 11413–11418 (2014). doi:10.1073/pnas.1411558111

37 Stracy, M. et al. Live-cell superresolution microscopy reveals the organization of RNA polymerase in the bacterial nucleoid. Proceedings of the National Academy of Sciences 112, E4390–E4399 (2015). doi:10.1073/pnas.1507592112

38 Mosteller, R. D. & Yanofsky, C. Transcription of the tryptophan operon in Escherichia coli: Rifampicin as an inhibitor of initiation. Journal of Molecular Biology 48, 525–531 (1970). 10.1016/0022-2836(70)90064-1

39 Binenbaum, Z., Parola, A. H., Zaritsky, A. & Fishov, I. Transcription- and translation-dependent changes in membrane dynamics in bacteria: testing the transertion model for domain formation. Molecular Microbiology 32, 1173–1182 (1999). 10.1046/j.1365-2958.1999.01426.x

40 Matsumoto, K., Hara, H., Fishov, I., Mileykovskaya, E. & Norris, V. The membrane: transertion as an organizing principle in membrane heterogeneity. Frontiers in Microbiology 6, 572 (2015). 10.3389/fmicb.2015.00572

41 Bellotto, N. et al. Dependence of diffusion in Escherichia coli cytoplasm on protein size, environmental conditions, and cell growth. eLife 11, e82654 (2022). 10.7554/eLife.82654

42 Linnik, D., Maslov, I., Punter, C. M. & Poolman, B. Dynamic structure of E. coli cytoplasm: supramolecular complexes and cell aging impact spatial distribution and mobility of proteins. Communications Biology 7, 508 (2024). 10.1038/s42003-024-06216-3

43 Chandran, V. & Luisi, B. F. Recognition of enolase in the Escherichia coli RNA degradosome. Journal of Molecular Biology 358, 8–15 (2006). 10.1016/j.jmb.2006.02.012

44 Chandran, V. et al. Recognition and cooperation between the ATP-dependent RNA helicase RhlB and ribonuclease RNase E. Journal of Molecular Biology 367, 113–132 (2007). 10.1016/j.jmb.2006.12.014

45 Nurmohamed, S., McKay, A. R., Robinson, C. V. & Luisi, B. F. Molecular recognition between Escherichia coli enolase and ribonuclease E. Acta Crystallographica Section D 66, 1036–1040 (2010). doi:10.1107/S0907444910030015

46 Nurmohamed, S., Vaidialingam, B., Callaghan, A. J. & Luisi, B. F. Crystal structure of Escherichia coli polynucleotide phosphorylase core bound to RNase E, RNA and manganese: implications for catalytic mechanism and RNA degradosome assembly. Journal of Molecular Biology 389, 17–33 (2009). 10.1016/j.jmb.2009.03.051

47 Stark, H. et al. The 70S Escherichia coli ribosome at 23 å resolution: fitting the ribosomal RNA. Structure 3, 815–821 (1995). 10.1016/S0969-2126(01)00216-7

48 Dai, X. et al. Reduction of translating ribosomes enables Escherichia coli to maintain elongation rates during slow growth. Nature Microbiology 2, 16231 (2016). 10.1038/nmicrobiol.2016.231

49 Miller, C. C. & Walker, J. The Stokes-Einstein law for diffusion in solution. *Proceedings of the Royal Society of London. Series A*, Containing Papers of a Mathematical and Physical Character 106, 724–749 (1924). doi:10.1098/rspa.1924.0100

50 Saffman, P. G. & Delbrück, M. Brownian motion in biological membranes. Proceedings of the National Academy of Sciences 72, 3111–3113 (1975). doi:10.1073/pnas.72.8.3111

51 Papanastasiou, M. et al. The Escherichia coli peripheral inner membrane proteome. Molecular & Cellular Proteomics 12, 599–610 (2013). 10.1074/mcp.M112.024711

52 Phillips, J. C. et al. Scalable molecular dynamics with NAMD. Journal of Computational Chemistry 26, 1781–1802 (2005). 10.1002/jcc.20289

53 Schuck, A., Diwa, A. & Belasco, J. G. RNase E autoregulates its synthesis in Escherichia coli by binding directly to a stem-loop in the rne 5′ untranslated region. Molecular Microbiology 72, 470–478 (2009). 10.1111/j.1365-2958.2009.06662.x

54 Apirion, D. Isolation, genetic mapping and some characterization of a mutation in Escherichia coli that affects the processing of ribonucleic acid. Genetics 90, 659–671 (1978). 10.1093/genetics/90.4.659

55 Bessarab, D. A., Kaberdin, V. R., Wei, C.-L., Liou, G.-G. & Lin-Chao, S. RNA components of Escherichia coli degradosome: evidence for rRNA decay. Proceedings of the National Academy of Sciences 95, 3157–3161 (1998). doi:10.1073/pnas.95.6.3157

56 Ghora, B. K. & Apirion, D. Structural analysis and in vitro processing to p5 rRNA of a 9S RNA molecule isolated from an rne mutant of E. coli. Cell 15, 1055–1066 (1978). 10.1016/0092-8674(78)90289-1

57 Li, Z., Pandit, S. & Deutscher, M. P. RNase G (CafA protein) and RNase E are both required for the 5′ maturation of 16S ribosomal RNA. The EMBO Journal 18, 2878–2885 (1999). 10.1093/emboj/18.10.2878

58 Roy, M. K., Singh, B., Ray, B. K. & Apirion, D. Maturation of 5-S rRNA: ribonuclease E cleavages and their dependence on precursor sequences. European Journal of Biochemistry 131, 119–127 (1983). 10.1111/j.1432-1033.1983.tb07238.x

59 Li, Z. & Deutscher, M. P. RNase E plays an essential role in the maturation of Escherichia coli tRNA precursors. RNA 8, 97–109 (2002).

60 Ow, M. C. & Kushner, S. R. Initiation of tRNA maturation by RNase E is essential for cell viability in E. coli. Genes & Development 16, 1102–1115 (2002). 10.1101/gad.983502

61 Ikeda, Y., Yagi, M., Morita, T. & Aiba, H. Hfq binding at RhlB-recognition region of RNase E is crucial for the rapid degradation of target mRNAs mediated by sRNAs in Escherichia coli. Molecular Microbiology 79, 419–432 (2011). 10.1111/j.1365-2958.2010.07454.x

62 Reyer, M. A. et al. Kinetic modeling reveals additional regulation at co-transcriptional level by post-transcriptional sRNA regulators. Cell Reports 36, 109764 (2021). 10.1016/j.celrep.2021.109764

63 Liou, G.-G., Jane, W.-N., Cohen, S. N., Lin, N.-S. & Lin-Chao, S. RNA degradosomes exist in vivo in Escherichia coli as multicomponent complexes associated with the cytoplasmic membrane via the N-terminal region of ribonuclease E. Proceedings of the National Academy of Sciences 98, 63–68 (2001). doi:10.1073/pnas.98.1.63

64 Szeto, T. H., Rowland, S. L., Habrukowich, C. L. & King, G. F. The MinD membrane targeting sequence is a transplantable lipid-binding helix. Journal of Biological Chemistry 278, 40050–40056 (2003). 10.1074/jbc.M306876200

65 Bayas, C. A. et al. Spatial organization and dynamics of RNase E and ribosomes in Caulobacter crescentus. Proceedings of the National Academy of Sciences 115, E3712–E3721 (2018). doi:10.1073/pnas.1721648115

66 Oviedo-Bocanegra, L. M., Hinrichs, R., Rotter, Daniel Andreas O., Dersch, S. & Graumann, P. L. Single molecule/particle tracking analysis program SMTracker 2.0 reveals different dynamics of proteins within the RNA degradosome complex in Bacillus subtilis. Nucleic Acids Research 49, e112–e112 (2021). 10.1093/nar/gkab696

67 Kapanidis, A. N., Uphoff, S. & Stracy, M. Understanding protein mobility in bacteria by tracking single molecules. Journal of Molecular Biology 430, 4443–4455 (2018). 10.1016/j.jmb.2018.05.002

68 Gray, W. T. et al. Nucleoid size scaling and intracellular organization of translation across bacteria. Cell 177, 1632–1648.e1620 (2019). 10.1016/j.cell.2019.05.017

69 Volkov, I. L. et al. tRNA tracking for direct measurements of protein synthesis kinetics in live cells. Nature Chemical Biology 14, 618–626 (2018). 10.1038/s41589-018-0063-y

70 Park, S. et al. Dynamic interactions between the RNA chaperone Hfq, small regulatory RNAs, and mRNAs in live bacterial cells. eLife 10, e64207 (2021). 10.7554/eLife.64207

71 Hamouche, L., Poljak, L. & Carpousis, A. J. Polyribosome-dependent clustering of membrane-anchored RNA degradosomes to form sites of mRNA degradation in Escherichia coli. mBio 12, e1932–1921 (2021). doi:10.1128/mbio.01932-21

72 Hamouche, L. et al. Dynamic membrane localization of RNase Y in Bacillus subtilis. mBio 11, e3337–3319 (2020). doi:10.1128/mbio.03337-19

73 Knight, J. D., Lerner, M. G., Marcano-Velázquez, J. G., Pastor, R. W. & Falke, Joseph J. Single molecule diffusion of membrane-bound proteins: window into lipid contacts and bilayer dynamics. Biophysical Journal 99, 2879–2887 (2010). 10.1016/j.bpj.2010.08.046

