## Supplemental Information for "Single-molecule imaging reveals the roles of the membrane-binding motif and the C-terminal domain of RNase E in its localization and diffusion in *Escherichia coli*"

### Supplementary Information

#### MATERIALS AND METHODS

##### Bacterial strains

Strains used in this study are listed in **Table S1**, and details of strain construction are in **Table S2**. All the engineered regions were confirmed by DNA sequencing.

Doubling times were measured from optical density at 600 nm ($OD_{600}$) using a microplate reader (Synergy HTX multi-mode reader, BioTek). Cultures were first grown overnight at 30°C and diluted $10^3$-$10^4$-fold in the same type of fresh media. Cultures were pipetted into a 96-well plate and grown for 24 hours at 30°C with continuous shaking and $OD_{600}$ measurements every 3 min.

The strains grown in M9 succinate were similarly grown overnight at 30°C but then diluted 100-fold in the same type of fresh media before growing for an additional ~24 hours. Cells were then diluted another 100-fold and pipetted into a 96-well plate for growth monitoring over 48 hours in the plate reader.

For data analysis, the background reading was removed by subtracting the initial $OD_{600}$ value, and the resulting values between 0.001 and 0.1 were fit to an exponential function to determine doubling time (listed in **Table S3**).

##### Sample preparation and cell growth for single-molecule imaging

Glass slides (Fisher 12-544-1) and #1.5 coverslips (Fisher 12544A or VWR 16004-344) were cleaned by three rounds of sonication in 100% ethanol, 70% ethanol, and Milli-Q water, respectively. They were stored in Milli-Q water until use and dried by nitrogen gas right before usage.

1% w/v agarose pads were created by melting agarose (Invitrogen 16500-100) in fresh liquid medium used for cell growth.

Unless otherwise stated, we grew cells in M9 minimal media supplemented with 0.2% v/v glycerol (Invitrogen 15514011), 0.1% w/v casamino acids (Bacto 223050), and 1 µg/mL thiamine (Research Products International T21020) at 30°C. A few colonies were inoculated in the M9 medium for at least 6 hours at 30°C with shaking at 220 rpm. This starter culture was diluted 3000fold or more for overnight growth. When the $OD_{600}$ reached 0.150-0.250, cells were placed on an agarose pad prepared on a glass slide. Excess liquid was removed by air drying for about a minute and then a cleaned coverslip was laid on the top and sealed with VALAP. The sample was imaged immediately. LacY, LacY2, LacY6, MTS, and RNE with *lacZ* overexpressed cells were induced with a final concentration of 1mM IPTG for at least 90 minutes or up to 16 hours before imaging.

**Drug treatment**

To see the effect of mRNA amount on the diffusion of proteins of interest, cells were treated with either rifampicin (400 µg/mL) for 15 min or chloramphenicol (100 µg/mL) for 30 min in the liquid culture with shaking at the growth temperature. Corresponding agarose pads for imaging included either 200 µg/mL rifampicin or 100 µg/mL chloramphenicol. We added these compounds when the molten agarose was at 55-65°C.

**Single-molecule imaging**

We took bright-field images and fluorescence images using a Nikon Ti-2 microscope equipped with a TIRF objective 100x/1.49 NA (Nikon), a 4-color laser launch (iChrome MLE-LFA-NI1), and an EM CCD camera (Andor iXon Ultra). For single-particle tracking photoactivated localization microscopy (sptPALM)[1], fluorescent images were taken using two lasers. A 405-nm laser was used to stochastically convert mEos3.2 from the green to the red fluorescence structure with a power between 0.003-0.055 W/cm$^2$ depending on the number of molecules of interest. A 561 nm laser was used to excite the photo-converted mEos3.2 molecules with a power between 2.3-2.7 W/cm$^2$. Images were taken with a frame rate of 21.7 ms with continuous laser illumination for 3 minutes. Each cell strain was imaged on at least three separate days, and 6-16 movies were collected each day.

All images were acquired using the Nikon Elements software (Nikon). We used a quasi-TIRF laser angle to reduce background. The same angle was used for all imaging.

**Nucleoid imaging**

We imaged the nucleoid region of cells containing RNE-mEos3.2 and HU-mCherry (SK512) grown in the same way as our single-molecule imaging samples. We used a Nikon Ti-2 microscope equipped with a phase-contrast Plan Apochromat objective 100x/1.45 NA (Nikon), a Sola SE II 365 light engine (Lumencor), an ET/mCh/TR filter cube (Nikon, 96365), and an Orca-R2 CCD camera (Hamamatsu Photonics). All images were acquired using the Nikon Elements software. Cells were treated with chloramphenicol in the same way as in single-molecule imaging of RNE-mEos3.2 (SK187). We also imaged cells without antibiotic treatment as a control.

**mRNA degradation rate measurement**

We measured mRNA levels and degradation rates as described previously[2]. Briefly, cells were grown the same way as for imaging experiments. When cells were in the early exponential phase ($OD_{600}$ ~0.02), cells were induced with 0.2 mM IPTG. The IPTG addition marks time zero (t = 0). At t = 75 s, cells were re-repressed with 500 mM glucose. During this time course, cells were sampled every 20-60 s for total RNA extraction using PureLink RNA Mini Kit (Life Technologies). Next, *lacZ* mRNA levels were quantified by qRT-PCR using KAPA SYBR FAST qPCR Master Mix (KAPA Biosystems) and CFX Connect Real-Time System (Bio-Rad). Primer sequences for 5' end and 3' end mRNA regions are provided in **Table S5**.

For the RNE over-expression strain (SK394) shown in **Fig. S7B**, cells were inoculated in the same way as other strains, and 0.2% arabinose was added at the time of dilution for overnight growth. When $OD_{600}$ became 0.1 the next day, cells were washed with fresh media of the same volume (with no arabinose), and the culture was shaken until $OD_{600}$ reached 0.2.

##### Cell fixation

To fix *E. coli* cells, we followed the protocol we have established previously[3]. Namely, cells were grown under the same conditions as used for microscopy. Once they reached the exponential phase, cells were fixed with 4% formaldehyde in 0.03 M disodium phosphate buffer (pH 7.4) for 15 minutes at room temperature, followed by an additional 30 minutes on ice. After fixation, cells were washed three times with PBS. For each wash, cells were pelleted by centrifugation at 5,000 × *g* for 4 minutes at room temperature. Cells were then resuspended in PBS and kept on ice until imaging. All samples were imaged on the same day as fixation.

#### DATA ANALYSIS

##### Single-molecule tracking

Data was analyzed in Matlab using Oufti[4] for cell detection from bright-field images, u-track[5] for fluorescent spot detection and linking into tracks, and a custom-built Matlab code called spotNorm for finding a fluorescent spot's normalized position in a cell (**Fig. 1A-C**). After this, the diffusion and subcellular localization were analyzed as described below.

*Oufti:* Cell outlines (or cell mesh) were computed from bright field images using Oufti software developed by Jacobs-Wagner Lab[4]. The following parameters were used:

edge mode for detection = valley, with dilate = 2
openNum = 1
InvertImage = 0
ThreshFactorM = 0.63493
ThreshMinLevel = 0.795
EdgeSigmaL = 1
EdgeSigmaV = 7
ValleyThresh1 = 0.14286
ValleyThresh2 = 0.77143

Only cells that are isolated (not touching other cells) and not dividing (based on cell constriction) were used in the analysis.

*u-track:* Fluorescent spots were detected and compiled into tracks using u-track software developed by Danuser Lab and Jaqaman Lab[5]. The parameters for fluorescent spot detection were as follows: pixel size of 160 nm, time interval of 0.0217s, numerical aperture of 1.49, and camera bit depth of 16 bits. Spot detection was done via Gaussian mixture-model fitting with the Gaussian standard deviation set to 0.9 pixels and the alpha-value for comparison with local background was set to 0.015. Also, the hypothesis test alpha value for Gaussian fitting at local maxima was set to an amplitude of 0.015 and distance of 0.05. For the tracking step, we used the default settings for 2D particle tracking with the maximum gap to close changed to 0 to avoid gaps in a track. Additionally, the minimum length of track segments from the first step was set to 4 frames. This setting ensures that all trajectories from u-track are at least 4 frames long.

A notable parameter in u-track is the max pixel linking distance. We set it to 1-3 pixels depending on the preliminary diffusion coefficient based on trajectories identified using u-track's default 5-pixel maximum linking distance, $D_{5pix}$. We found that the max linking distance of 5 pixels identifies artificial large random jumps in our data set that falsely creates a large diffusion coefficient. To avoid this error, we set the max linking distance radius as ceiling(2*sqrt(4$D_{5pix}\Delta t$)), where $\Delta t$ is the frame time (0.0217 s). Alternatively, we can use 1 pixel if $D_{5pix} < 0.0737$ μm$^2$/s and 2 pixels if 0.0737 μm$^2$/s $< D_{5pix} <$ 0.295 μm$^2$/s.

*spotNorm:* Fluorescent spot positions in (x,y) pixel coordinates, obtained from u-track, were transformed into normalized coordinates along the cell's long axis (yNorm) and short axis (xNorm) using a custom-built MATLAB script called spotNorm. First, the fluorescent spot position (in pixels) was compared to the cell outlines (in pixels) from Oufti of all cells for the corresponding fluorescent movie. Only spots within a cell were used in the downstream analysis. The center of the cell outline and the centroid region of spots inside of the cell were compared to check for drift between taking the bright-field image and the start of fluorescence tracking. If the centroid regions were different, then the average distance between the cell outline and the fluorescent data of all the cells in the frame were used to calculate a correction shift for fluorescent data. The spot's position was then projected onto the long and short axes of the cell, and the projected distances were normalized by the corresponding cell axis lengths. A positive or negative value for the spot's xNorm position was determined by the sign of the cross-product of the spot within the center line of the cell. xNorm values were between -1 to 1. yNorm values were between 0 and 1 where 0 and 1 are the two different cell poles.

By combining xNorm data from many cells, we created xNorm histograms. Spots included in these histograms were only from the cylindrical portion of the cell, not from the hemispherical endcaps. This was calculated based on the spot position along the long axis, y, being within R < y < (L-R), where R is half of the mean cell width and L is the length of the cell. For better statistics, we included spots from the first frames of a track in the xNorm histogram. Because we set a minimum track length of 4 frames in u-track analysis, each track contributes 4 points equally to the xNorm histogram. The bin size for xNorm histograms is 0.04. In the xNorm histogram, the error was calculated from the SEM from bootstrapping. Each xNorm histogram is displayed with SEM-based error ranges by a shaded area, but it is smaller than the data line width. RNE's xNorm histogram error shaded region is compared to error bar lines (**Fig. S1A**).

We used a *D* cutoff to examine xNorm of the fast and slow trajectories separately (**Fig. S2**). The $D_{cutoff}$ was chosen to match the fraction of the slow population ($D<D_{cutoff}$) with the MB% of that strain. The xNorm for the slow populations typically had LacY-like xNorm profile, whereas the fast population had one flat central peak, similar to the distribution of RNE ΔMTS (**Fig. S2**). Note that only tracks with at least 12 frames will have a calculated *D*. Thus, only a subset of the xNorm data was used in the $D_{cutoff}$ xNorm histograms.

Diffusion: Tracks lasting at least 12 frames long were used for diffusion analysis. Time-averaged MSD was calculated according to eq. 1, and the diffusion coefficient (*D*) was calculated by a linear fit using eq. 2[6]:

$$MSD(\tau) = \frac{1}{N_\tau}\sum_{i=1}^{N_\tau}[\vec{r}(t_i) - \vec{r}(t_i+\tau)]^2 \quad (1)$$

$$MSD(\tau) = 4D\tau + b, \text{where } b = -\frac{4D\Delta t}{3} + 4\sigma^2 \quad (2)$$

In eq. 1, $\vec{r}(t_i)$ is the 2D vector location of a particle at time $t_i$, $\tau$ is the lag time, and $N_\tau$ is the number of frames that are averaged over in a track for that lag time. In eq. 2, $\Delta t$ is the exposure time, and $\sigma$ is the localization error. The y-intercept, *b*, is composed of two terms. The first term is a correction for dynamic error, and the second term is for static error[6]. We only fit the first three time points of MSD with eq. 2 to get *D*. This ensures that the fit is done on short time delays (the first 25% of the minimum track length) to avoid MSD at large time intervals with low statistics[7,8]. Once we obtained *D* from individual tracks, we calculated their mean and SEM (standard deviation divided by the square root of the sample size) to report mean ± SEM of the diffusion coefficient.

We checked slow and fast subpopulations of proteins by analyzing the distribution of *D* of individual tracks. First, we fit the distribution of $D_{RNE}$ using a single Gaussian distribution (eq. 3) or a two-population distribution (eq. 4) (**Fig. S3B-C)**. We assumed MB% for the slow population and the rest for the fast population in the two-population fitting (**Fig. S3C**). The residuals from a one-population fit and a two-population fit were similar for RNE, indicating that the single Gaussian fit can be used. The distribution of *D* of ribosomal proteins (L1) showed a long tail, suggesting more than one mobility population exists, in agreement with other studies reporting multiple mobility populations for ribosomal proteins[9,10]. Hence, we used a two-population Gaussian distribution (eq. 4) for fitting (**Fig. S3B**). The slow population is likely from polysomes, or the ribosomes bound to mRNA (as these form a larger complex), and the fast population may represent free subunits not engaged in translation.

$$g(D_h) = \frac{c}{\sigma\sqrt{2\pi}} \exp\left(-\frac{1}{2}\frac{(D_h-\mu)^2}{\sigma^2}\right) \tag{3}$$

$$g(D_h) = \frac{c}{\sqrt{2\pi}}\left[\frac{a}{\sigma_1}\exp\left(-\frac{1}{2}\frac{(D_h-\mu_1)^2}{{\sigma_1}^2}\right) + \frac{(1-a)}{\sigma_2}\exp\left(-\frac{1}{2}\frac{(D_h-\mu_2)^2}{{\sigma_2}^2}\right)\right] \tag{4}$$

In eq. 3, $c$ is the scaling factor, $D_h$ is the diffusion coefficients from $D$ histograms, $\mu$ is the mean, and $\sigma$ is the standard deviation. Similarly, in eq. 4, $\mu_1$ and $\mu_2$ are the means of the first and second populations, $\sigma_1$ and $\sigma_2$ are the standard deviations of the first and second populations, $a$ is the fraction of the first population, and (1-$a$) is the fraction of the second population. As mean values are for diffusion coefficients, we also refer to $\mu$ as $D_1$ for the single Gaussian fitting and $\mu_1$ and $\mu_2$ as $D_1$ and $D_2$, respectively, for the two-population Gaussian fitting.

**Membrane-binding percentage (MB%)**

To model an xNorm histogram from a distribution of molecules in 3D space, we assume that a cell is a cylinder, where molecules are homogeneously distributed on the surface (membrane) or inside (cytoplasm). We use a Cartesian coordinate, in which the long axis of the cylinder is aligned along the y-axis with its circular cross-section oriented vertically in the x-z plane (**Fig. S1D**).

Since molecules are uniformly distributed, we mainly consider the distribution of molecules in the circular cross-section (perpendicular to the y-axis) and calculate a marginal distribution of the $x$-coordinates, as $P_{memb}$ and $P_{cyto}$ for molecules on the circumference and inside, respectively. Finally, the probability distribution is modeled as a mixture distribution of the molecules on the membrane and in the cytoplasm.

$$P(x) = m \cdot P_{memb}(x) + (1-m) \cdot P_{cyto}(x) \tag{5}$$

where $m$ is the membrane-binding percentage (MB%).

In detail, the inner membrane is described as a circle with radius $r$. Define $z(x) = \sqrt{r^2 - x^2}$. Then, the conditional probability distributions for molecules on the membrane and in the cytoplasm are as follows:

$$dP^0_{memb} = \frac{dx}{\pi z} \tag{6}$$

$$dP_{cyto}^{0} = \frac{2z \cdot dx}{\pi r^2} \quad (7)$$

Next, we introduce imaging effects to the model as follows: the localization error was added by convolving the raw distribution with a 2D-Gaussian blur $g(x,z) = \phi(x,0,\sigma) \cdot \phi(z,0,\sigma)$, where $\phi(u,c,s) = e^{-(u-c)^2/(2 \cdot s^2)}/\sqrt{2\pi}s$ is a Gaussian kernel. Here, we assumed the same localization uncertainty (σ) in the x and z dimensions. While we could not justify this assumption, we found that the choice of uncertainty in the z dimension minimally affects the final outcome (xNorm histogram), and hence the uncertainty in the x dimension was used to simplify the model. Additionally, the imaging system has a certain depth of focus, beyond which the signal cannot be captured by the camera. We let the defocused range at the bottom part of the cell be given by $z < -f$ (**Fig. S1D**). With these adjustments, the molecular density along the $z$-axis is proportional to $\int_{-\infty}^{f} \phi(u,-z,\sigma)du = \Phi((f+z)/\sigma)$, where Φ is the cumulative distribution function of a standard Gaussian. Therefore, the conditional probability distributions are formulated as:

$$dP_{memb} \propto \left(\Phi\left(\frac{f+z}{\sigma}\right) + \Phi\left(\frac{f-z}{\sigma}\right)\right) \cdot dP_{memb}^{0} * g \quad (8)$$

$$dP_{cyto} \propto \left(\int_{-z}^{z} \Phi\left(\frac{f+z}{\sigma}\right) du\right) \cdot \frac{dP_{cyto}^{0}}{z} * g \quad (9)$$

Here, ∝ simply means the density should be scaled by normalizing constants.

To fit the experimental data to the model, parameters need to be scaled. The inner membrane radius $r$ is $R/dilF$, where $R$ is half of the cell width (experimentally obtained from bright-field images and Oufti analysis) and $dilF$ is the dilation factor, which accounts for the fact that membrane-bound molecules (from mEos3.2. imaging) are localized away from the cell boundary identified by the bright-field images (e.g., **Fig. 1C**). In the end, $R$ is normalized to 1 to provide xNorm. Also, $f = R - fCut$ (**Fig. S1D**), and hence, $fCut$ is zero when the view of the cell is unobstructed. $\sigma = locErr/1000$, the localization error measured in μm. Theoretical xNorm distributions of membrane and cytoplasmic molecules are illustrated in **Fig. S1E**-**G**.

To estimate the MB% and other parameters in experimental xNorm data, we used the Markov-Chain Monte Carlo algorithm, which is an effective simulation approach for estimating Bayesian posterior distributions of multi-parameter models and their representative statistics. For better statistics, we fitted the absolute xNorm histogram. Specifically, we constructed a histogram by binning the experimental data (absolute xNorm from 0 to 1) and minimizing the mean-squared error to the model-implied values. All parameters ($dilF$, $locErr$, $fCut$, $m$) were free to change within the intuitive bounds (e.g. $m \in [0,1]$), and they have uniform priors. We implemented the fitting procedure in

Python using the *pymcmcstat* package[11]. For robustness, we examined different choices of hyper-parameters (bin size, bounds, sampling numbers, and the burn-in period) and checked that the end result is insensitive to them. The fitting results are provided in **Table S4**.

##### mRNA degradation rate

We estimated mRNA degradation rates by fitting a single-exponential decay model to the Z5 time-course data, as described previously[2]. Briefly, the exponential fits were performed between t = ~150 s to ~210 s for $k_{d1}$ and between t = ~300 s to 600 s for $k_{d2}$.

##### Statistical test

We tested for statistical similarity using *p*-values determined by a two-tailed Student's *t*-test on small data sets for *lacZ* mRNA degradation (**Table S7**). The two-tailed Student's *t*-test was calculated using the ttest2 function in MATLAB at the 5% significance level. When the alternative hypothesis was checked for differences, no additional specifications were used. When the alternative hypothesis was for the mean to be less than the other, we used the additional specification "tail", "left". If the *p*-value is above 0.05, the two data sets are statistically similar. Otherwise, the alternative hypothesis is supported.

##### Detection efficiency of cytoplasmic vs membrane-bound proteins

Since proteins diffuse more rapidly in the cytoplasm than in the membrane, they might not be detected as well as the membrane-bound proteins in our microscopy experiments. To assess this potential undercounting, we compared the number of tracks in live vs fixed cells for WT RNE and for cytoplasmic RNE ΔMTS. Fixed cells were imaged as previously described, except that the agarose pad was made of PBS instead of growth media. U-track was used to find the number of tracks in the cell using a max linking distance of 1 pixel. The mean number of tracks per cell in live and fixed cells were compared using the Fisher test (**Fig. S1C**). The Fisher-Irwin test is a statistical significance test used for contingency tables. The returned p-value was 0.18, indicating insignificant bias introduced by localization (and mobility) in detection.

##### Cluster analysis

We analyzed clustering of RNE based on mEos3.2 images acquired from fixed cells, in which clustering is preserved. We used spatial point statistics used in a previous study of RNE clustering in *C. crescentus*[12]. For the analysis, we first chose cells with at least 300 spots (or tracks) located throughout the cell. A complete spatial randomness (CSR) was simulated as molecules homogeneously distributed on the membrane of a 3D cell using the same length, width, and number of points as each imaged cell. The simulated spots were projected to a 2D plane for comparison with experimental data.

Global clustering was quantified using Ripley's K function[13] and its normalized forms[14]. Ripley's K function represents the average probability of finding localizations at a distance r from the typical localization[15]. For CSR, K(r) scales with the cell volume and is described by Eq. 10, where $b_d r^d$ is the volume of the unit sphere in d dimensions. Deviations from this shape indicate clustering or repulsion. The normalized K-function is given by L(r) in Eq. 11, which can further be normalized to the H(r) function[16]. For example, the expected value of K(r) for a random Poisson distribution is $\pi \cdot r^2$, and L(r) = r, H(r) = 0.

To analyze clustering across many cells, $\int_r H_{data}(r) - H_{CSR}(r)$ was calculated for each cell.

$$K(r) = b_d r^d \tag{10}$$

$$L(r) = \sqrt[d]{K(r)/b_d} \tag{11}$$

$$H(r) = L(r) - r \tag{12}$$

**Code availability**

MATLAB code for data analysis and Python code for modeling are available at the publicly accessible site, https://github.com/sjkimlab/Code_Publication.

#### ALL-ATOM MOLECULAR DYNAMICS SIMULATIONS

The all-atom simulations were performed using the NAMD 3.0 software[17]. The CHARMM-GUI Membrane Builder[18-20] was utilized to generate *E. coli* lipid bilayers consisting of 80 POPE and 20 POPG lipids per leaflet. The lipid membranes were hydrated using the TIP3P water model. The protein-membrane system was equilibrated with the CHARMM36 force field[21]. Simulations were conducted under the NPT ensemble, maintaining the temperature at 310 K and pressure at 1 bar using the Nosé-Hoover Langevin piston and Langevin dynamics. A 2-fs time step was employed for integrating the equations of motion. van der Waals and electrostatic interactions were truncated at a 12 Å cutoff, with a switching function applied between 10 and 12 Å. The Particle-Mesh Ewald (PME) method was used to handle long-range electrostatic interactions. For MTS, we used the 17 amino acid sequence: QPGLLSRFFGALKALFS. For LacY2, we used the 74 amino acid sequence: MYYLKNTNFWMFGLFFFFYFFIMGAYFPFFPIWLHDINHISK SDTGIIFAAISLFSLLFQPLFGLLSDKLGLRK. For LacY6, we used first 193 residues of LacY. For LacY12, we used the full-length LacY (417 residues).

We estimated diffusion by tracking the center-of-mass positions of the proteins over time. Ensemble-averaged MSD was calculated by averaging the squared displacements from 5 replicas (**Fig. S4A**). The diffusion coefficient was then obtained based on the MSD = 4*D*t relationship.

To explain the unexpected diffusion dynamics of MTS relative to LacY2, we calculated the per-residue interaction energies between the protein and membrane systems using the NAMDEnergy plugin in VMD (**Fig. S4B**). The interaction energy per residue for MTS is more negative compared to LacY2, indicating a stronger affinity for the lipid head groups. In contrast, LacY2 is primarily situated within the hydrophobic tails of the membrane lipids, showing a weaker interaction.

#### SUPPLEMENTARY DISCUSSION

##### Minimum diffusion coefficient that can be measured using our microscope

To test the minimum *D* that our microscope can measure for mEos3.2, we prepared surface-immobilized mEos3.2 and measured *D* of the stationary molecules. We purified His6-streptavidin-mEos3.2 expressed from a plasmid (SK567) in BL21 cells using His-Spin Protein Miniprep (Zymo Research, P2001) and then buffer exchanged into PBS (pH 7.4) using a centrifugal filter unit (Merck Millipore, UFC510024). His6-streptavidin-mEos3.2 were pipetted onto a coverslip functionalized with biotin and allowed to bind for 5 min before washing away excess liquid. After washing with Milli-Q water, it was sealed with a cleaned glass slide. Surface-immobilized mEos3.2 molecules were imaged the same way we imaged mEos3.2 in *E. coli*. The diffusion of His6-streptavidin-mEos3.2 immobilized on biotin coverslips was 0.0020 ± 0.0001 $\mu m^2/s$ (**Fig. S3A**). This determines to what extent experimental noise between the fluorophore and microscope's localization error can cause apparent diffusion of a stationary molecule. This *D* value was well below our measured diffusion coefficients of the molecules in this study, indicating slow diffusion coefficients are not due to tracking limitations.

##### Calculation of the mass of the RNA degradosome

The canonical RNA degradosome in *E. coli* is comprised of RNE, enolase, PNPase, and RhlB[22,23]. However, there are other proteins that have been shown to associate with the degradosome, such as Ppk, CsdA, and PAP[24-26].

We calculated the mass of the RNA degradosome assuming a fully loaded canonical complex, built upon RNE tetramer[27], in which each RNE monomer interacts with one RhlB dimer, one enolase dimer, and one PNPase trimer. According to the amino acid sequence or reported mass in the literature (for enolase[28]), the masses are as follows.

RNE monomer = 118 kDa
RhlB monomer = 47.1 kDa
Enolase monomer = 45.7 kDa
PNPase monomer = 77.1 kDa

Considering a linker (5-7 amino acids depending on the strain, or 0.56-0.78 kDa) and mEos3.2 (27.2 kDa based on the sequence), we estimate the effective mass of WT RNE-mEos3.2 complex to be ~2.27 MDa. Other strains used in this study (such as LacY variants or MTS) were calculated based on the amino acid sequence including a linker and mEos3.2.

##### Effect of mEos3.2 fusion to RNE function

We checked the effect of a fluorescent protein attached to RNE on *lacZ* mRNA degradation rates. Both $k_{d1}$ and $k_{d2}$ are similar in cell strains containing RNE with or without mEos3.2 attached (**Fig. S7A**). Additionally, the fluorescent protein did not change the doubling times for these strains (**Table S3**).

##### Effect of chloramphenicol treatment and induction of *lacZ* mRNA from plasmids

The diffusion coefficient of RNE was unaffected by chloramphenicol treatment or overexpression of *lacZ* mRNA (**Fig. S3F**). To support our observation, we tested if the chloramphenicol treatment resulted in nucleoid compaction due to an increase in polysome concentrations—an effect shown in previous studies[29,30]. As shown in **Fig. S3D**, SK512 (RNE-mEos3.2, HU-mCherry) cells showed smaller nucleoid upon chloramphenicol addition. Furthermore, we tested if *lacZ* mRNAs were indeed over-expressed by measuring the mRNA levels relative to a reference gene, *gapA* by RT PCR. The IPTG induced cells (SK411) showed about 25 times higher *lacZ* mRNA levels than those from uninduced cells (**Fig. S3E**), supporting that the overexpression was effective.

#### SUPPLEMENTARY TABLES

##### Table S1: List of strains used in this study

| Strain number | Genotype | Source | Use |
|---|---|---|---|
| SK1 | MG1655 | | Cloning<br>Doubling time |
| SK47 | BW25993 *rplA::rplA-mEos2* | J. Elf[10] | Imaging |
| SK52 | MG1655 Δ*araFGH araE*::P13-*araE* | C. Jacobs-Wagner | Cloning |
| SK72 | NCM3416 *rne::rne-mCherry* FRT-*cat*-FRT | A. Carpousis[31] | Cloning |
| SK98 | MG1655 Δ*lacYA* | C. Jacobs-Wagner, CJW5461[32] | qRT-PCR<br>Cloning |

| | | | |
|---|---|---|---|
| SK105 | MG1655 Δ*lacIZYA* | C. Jacobs-Wagner, CJW6643[32] | Cloning |
| SK107 | MG1655 *rne::rne*(ΔMTS)-*mCherry* | C. Jacobs-Wagner, CJW5685[33] | Cloning |
| SK186 | MG1655 *rne::rne*(1-592)-*yfp* FRT-*kan*-FRT | C. Jacobs-Wagner, JRH474 | Cloning |
| SK187 | MG1655 *rne::rne-mEos3.2* FRT-*kan*-FRT | C. Jacobs-Wagner, JRH475 | Imaging |
| SK213 | BW25113 *hupA::hupA-mcherry* FRT-*kan*-FRT | C. Jacobs-Wagner, CJW5158[19] | Cloning |
| SK249 | MG1655 *rne::rne*ΔMTS-*mEos3.2* FRT-*kan*-FRT | This study | Imaging |
| SK290 | MG1655 *rne::rne-mEOS3.2* | This study | Cloning |
| SK292 | MG1655 *lacYA::lacY-mEos3.2* FRT-*kan*-FRT | This study | Imaging |
| SK304 | MG1655 *rne::rne-mEos3.2, ΔrhlB::*FRT-*kan*-FRT | This study | Imaging |
| SK308 | MG1655 *rne::rne-mEos3.2, Δpnp::*FRT-*kan*-FRT | This study | Imaging |
| SK360 | MG1655 *ΔaraFGH araE*::P13-*araE araBAD*::*rne-yfp-kan rne*::*rne-mcherry* FRT-*cat*-FRT | This study | Cloning |
| SK364 | MG1655 *ΔaraFGH araE*::P13-*araE araBAD*::*rne-yfp rne*::*rne-mcherry* | This study | Cloning |
| SK370 | MG1655 Δ(*lacYA*) *rne::rne*(1-592)-*yfp* FRT-*kan*-FRT | Our lab[2] | qRT-PCR |
| SK373 | MG1655 *rne::rne*(1-529)-*mEos3.2* FRT-*kan*-FRT | This study | Imaging |
| SK374 | MG1655 *rne::rne*(1-592)-*mEos3.2* FRT-*kan*-FRT | This study | Imaging |
| SK384 | MG1655 *rne::rne*(1-592)-*yfp* | This study | Cloning |
| SK394 | MG1655 Δ*lacYA ΔaraFGH araE*::P13-*araE araBAD::rne-yfp rne::rne-mcherry* | This study | qRT-PCR |
| SK404 | MG1655 *rne::rne*(1-564)-*lacY-mEos3.2* FRT-*kan*-FRT | This study | Imaging |
| SK405 | MG1655 Δ*(lacYA) rne::rne*(1-564)-*lacY-mEos3.2* FRT-*kan*-FRT | This study | qRT-PCR |
| SK407 | MG1655 *lacZYA::lacZ-mEos3.2* FRT-*kan*-FRT | This study | Imaging |
| SK411 | MG1655 *rne::rne-mEos3.2*<br>pUC19-lacI-lacZonly (amp) | This study | Imaging |
| SK424 | MG1655 *lacYA::lacY*(1-73)-*mEos3.2* FRT-*kan*-FRT | This study | Imaging |
| SK425 | MG1655 *lacYA::lacY*(1-192)-*mEos3.2* FRT-*kan*-FRT | This study | Imaging |

| | | | |
|---|---|---|---|
| SK455 | MG1655 *ΔlacIZYA* pUC19-lacI-plac-mEos3.2-MTS (amp) | This study | Imaging |
| SK466 | MG1655 *rne::rne*(1-564)-*lacY*(1-73)-*rne*CTD-*mEos3.2* FRT-*kan*-FRT | This study | Imaging |
| SK467 | MG1655 *rne::rne*(1-564)-*lacY*(1-192)-*rne*CTD-*mEos3.2* FRT-*kan*-FRT | This study | Imaging |
| SK482 | MG1655 *rne*::*rne-venus hupA*::*hupA-mcherry* FRT-*kan*-FRT | This study | Imaging |
| SK486 | MG1655 *rne*::*rne*(1-592)-*venus hupA*::*hupA-mcherry* FRT-*kan*-FRT | This study | Imaging |
| SK505 | MG1655 Δ(*lacYA*) *rne::rne*(1-564)-*lacY*(1-73)-*rne*CTD-*mEos3.2* FRT-*kan*-FRT | This study | qRT-PCR |
| SK506 | MG1655 Δ(*lacYA*) *rne::rne*(1-564)-*lacY*(1-192)-*rne*CTD-*mEos3.2* FRT-*kan*-FRT | This study | qRT-PCR |
| SK507 | MG1655 *rne::rne*(1-564)-*lacY*(1-73)-*mEos3.2* FRT-*kan*-FRT | This study | Imaging |
| SK508 | MG1655 Δ(*lacYA*) *rne::rne*(1-564)-*lacY*(1-73)-*mEos3.2* FRT-*kan*-FRT | This study | qRT-PCR |
| SK512 | MG1655 *rne::rne-mEos3.2 hupA::hupA-mcherry kan* | This study | Imaging |
| SK592 | MG1655 *rne::rne*(1-564)-*lacY*(1-192)-*mEos3.2* FRT-*kan*-FRT | This study | Imaging |
| SK593 | MG1655 Δ(*lacYA*) *rne::rne*(1-564)-*lacY*(1-192)-*mEos3.2* FRT-*kan*-FRT | This study | qRT-PCR |
| SK594 | MG1655 Δ(*lacYA*) *rne::rne*(1-592)-*yfp* | This study | Cloning |
| SK595 | MG1655 Δ(*lacYA*) *rne::rne-mEos3.2* FRT-*kan*-FRT | This study | qRT-PCR |
| SK598 | MG1655 Δ(*lacYA*) *rne::rne*(1-564)-*lacY-rne*CTD-*mEos3.2* FRT-*kan*-FRT | This study | Imaging qRT-PCR |
| SK741 | MG1655 Δ(*lacYA*) *rne*::*rne*(1-564)-MTS(F574AF575A)- *rne*CTD -*mEos3.2* FRT-*kan*-FRT | This study | Imaging qRT-PCR |
| SK742 | MG1655 Δ(*lacYA*) *rne*::*rne*(1-564)-MTS(F575E)-*rne*CTD-*mEos3.2* FRT-*kan*-FRT | This study | Imaging qRT-PCR |
| SK743 | MG1655 Δ(*lacYA*) *rne*::*rne*(1-564)-MTS(F582E)-*rne*CTD-*mEos3.2* FRT-*kan*-FRT | This study | Imaging qRT-PCR |
| SK748 | MG1655 Δ(*lacYA*) *rne*::*rne*(1-564)-MTS(F574AF575A)-*mEos3.2* FRT-*kan*-FRT | This study | Imaging qRT-PCR |
| SK749 | MG1655 Δ(*lacYA*) *rne*::*rne*(1-564)-MTS(F575E)- | This study | Imaging |

| | | | |
|---|---|---|---|
| | *mEos3.2* FRT-*kan*-FRT | | qRT-PCR |
| SK750 | MG1655 Δ(*lacYA*) *rne*::*rne*(1-564)-MTS(F582E)-*mEos3.2* FRT-*kan*-FRT | This study | Imaging qRT-PCR |
| **Plasmids** | | | |
| pBAD18Kan | | [34] | Cloning |
| pET29b-H6_Streptavidin_sfGFP | | A gift from Mark Arbing (Addgene plasmid # 124296; http://n2t.net/addgene:124296; RRID: Addgene_124296) | Cloning |
| pKD13 | | [35] | Cloning |
| pUC19 | | | Cloning |
| SJK1606 | pBAD18kan-mEos3.2-MTS | This study | Cloning |
| SJK1689 | pUC19-lacI-lacY2-CTD-mEos3.2-Kan | This study | Cloning |
| SJK1697 | pUC19-lacI-lacY6-CTD-mEos3.2-frtKanfrt | This study | Cloning |
| SJK1716 | pUC19-lacI-lacY12-CTD-mEos3.2-frtKanfrt | This study | Cloning |
| SK141 | pUC19-lacI-lacZonly | C. Jacobs-Wagner, CJW6647[32] | Cloning |
| SK189 | pBAD18 rne-yfp-kan | C. Jacobs-Wagner, JRH515 | Cloning |
| SK567 | pET29b-H6_Streptavidin_mEos3.2 | This study | Cloning |

#### Table S2: Strain construction

| Strain number | Construction |
|---|---|
| SK187 | *mEos3.2-kan* was integrated to the end of *rne* on the chromosome of MG1655 by lambda Red recombination. |
| SK189 | *rne-yfp* sequence was from pVK207[36]. |
| SK249 | *mEos3.2-kan* region in SK187 was amplified using the following primers and integrated into SK107 to replace *mCherry* at the end of *rne*ΔMTS by lambda Red recombination.<br>SJK033: GTGCCGCAGGTGGTCATACG<br>SJK034: GGTTAGCAAGGATGCCATTCG |
| SK290 | The *kan* cassette in SK187 was removed by FLP recombination. |

| | |
|---|---|
| SK292 | *mEos3.2-kan* region in SK187 was amplified using the following primers and integrated to the end of *lacY* in MG1655 by lambda Red recombination.<br>K018:<br>GCGGCCCCGGCCCGCTTTCCCTGCTGCGTCGTCAGGTGAATGAAGTCGCTAGAGGTGGTTTATCCATGTCGGCGATCAAGCCGGAC<br>K019:<br>GCTGAACTTGTAGGCCTGATAAGCGCAGCGTATCAGGCAATTTTTATAATTTATCCTTAGTTCCTATTCC |
| SK304 | The *kan* cassette was amplified from pKD13 using the following primers and integrated into the chromosome of SK290 to replace *rhlB* gene by lambda Red recombination.<br>rhlB_KO_F:<br>CGGATACGCTTTCGTAAAGCAATAGTAAGCTGATATTCTACCACACTATGATTCCGGGGATCCGTCGACC<br>rhlB_KO_R:<br>TGAATGATTTTGAGTATGACATTTTTTATTTAACCTGAACGACGACGATTTGTAGGCTGGAGCTGCTTCG |
| SK308 | The *kan* cassette was amplified from pKD13 using the following primers and integrated into the chromosome of SK290 to replace *pnp* gene by lambda Red recombination.<br>pnp_KO_F:<br>CCCGCCGCAGCGGAGGGCAAATGGCAACCTTACTCGCCCTGTTCAGCAGCATTCCGGGGATCCGTCGACC<br>pnp_KO_R:<br>ACACCAGTGCCGTAAGGTACTGTCTAAGAAAGAGAAAGGATATTACATTGTGTAGGCTGGAGCTGCTTCG |
| SK360 | First, *rne-mcherry-cat* in SK72 was moved to SK52 via phage transduction (SK349). Next, *rne-yfp-kan* was amplified from SK189 using primer K050 and K053 and integrated into *araBAD* locus on the chromosome of SK349 by lambda Red recombination.<br>K050:<br>GCAACTCTCTACTGTTTCTCCATACCCGTTTTTTTGGATGGAGTGAAACGATGAAAAGAATGTTAAT<br>K053:<br>GCTTGAGTATAGCCTGGTTTCGTTTGATTGGCTGTGGTTTTATACAGTCAAAGTATATATGAGTAAACTTGG |
| SK364 | From SK360, both *kan* and *cat* cassettes were removed by FLP recombination. |
| SK373 | *mEos3.2-kan* region in SK187 was amplified using the primers K066 and SJK034 and then integrated into the *rne* region in MG1655 by lambda Red recombination.<br>K066:<br>CGTCTGAAGAAGAGTTCGCTGAACGTAAGCGTCCGGAACAACCTGCGCTGCTCGAGGGTCCGGCTGGTCTGATGTCG |
| SK374 | *mEos3.2-kan* region in SK187 was amplified using the primers K065 and SJK034 and then integrated into MG1655 by lambda Red recombination.<br>K065:<br>GCGCACTGAAAGCGCTGTTCAGCGGTGGTGAAGAAACCAAACCGACCGAGCTCGAGGGTCCGGCTGGTCTGATGTCG |
| SK384 | The *kan* cassette in SK186 was removed by FLP. |

| | |
|---|---|
| SK394 | To make a clean *lacYA* deletion, *cat-sacB* from pEL04 was integrated into *lacYA* region in SK364 and then replaced by synthetic DNA lacZAfor and its complementary lacZArev.<br>lacZAfor:<br>AGCTGAGCGCCGGTCGCTACCATTACCAGTTGGTCTGGTGTCAAAAATAAATTA TAAAAATTGCCTGATACGCTGCGCTTATCAGGCCTACAAGTTCAGC |
| SK404 | *lacY-mEos3.2-kan* region was amplified from SK292 using primer K088 and K089 and used to replace the second half of rne in the chromosome of MG1655 by lambda Red recombination.<br>K088:<br>CGCCTGTTGTAGCTCCAGCACCGAAAGCTGCACCGGCAACACCAGCAGCTTAC TATTTAAAAAACACAAACTTTTGG<br>K089:<br>AATAAAAAAGCCCTGGCAGTTACCAGGGCTTGATTACTTTGAGCTAATTATTATC CTTAGTTCCTATTCC |
| SK405 | Same as SK404, but the DNA fragment was integrated into SK98. |
| SK407 | *mEos3.2-kan* region was amplified from SK292 using primer K098 and lacA_out50R and integrated into the end of *lacZ* in MG1655 by lambda Red recombination.<br>K098:<br>TCCAGCTGAGCGCCGGTCGCTACCATTACCAGTTGGTCTGGTGTCAAAAAAGA GGTGGTTTATCCATGTCG<br>lacA_out50R: GCTGAACTTGTAGGCCTGATAAGC |
| SK411 | SK141 plasmid was electroporated into SK290. |
| SK424 | *mEos3.2-kan* was amplified from SK187 using primer K099 and K028 and integrated into the *lacY* region in MG1655 by lambda Red recombination.<br>K099:<br>TATTCCAACCGCTGTTTGGTCTGCTTTCTGACAAACTCGGGCTGCGCAAAAGAG GTGGTTTATCCATGTCGGCGATCAAGCCGGAC<br>K028:<br>TGTAGATCGCTGAACTTGTAGGCCTGATAAGCGCAGCGTATCAGGCAATTTATC GTGAGGATGCGTCATCG |
| SK425 | Similar to SK424, but K101 and K028 primers were used to prepare the DNA fragment.<br>K101:<br>CACTCATCCTCGCCGTTTTACTCTTTTTCGCCAAAACGGATGCGCCCTCTAGAG GTGGTTTATCCATGTCGGCGATCAAGCCGGAC |
| SK455 | The plasmid was made by Gibson ligation of two fragments: (1) pUC19 backbone and *lacI* region of plasmid SK141 using two primers: lacZp_rev and lacA_out20F and (2) *mEos3.2*-MTS from SJK1606 (3'MTS). Here, MTS is a 51 base sequence from *rne*, and *mEos3.2* sequence is fused at the 5' side. The resulting plasmid was electroporated into SK105.<br>lacZp_rev: CATAGCTGTTTCCTGTGTGAAATTGTTATCC<br>lacA_out20F: ATTATAAAAATTGCCTGATACG |
| SK466 | *lacY*-CTD-*mEos3.2-kan* was amplified from plasmid SJK1689 using K088 and K089 and integrated into the *rne* region in SK384 by lambda Red recombination. |
| SK467 | Same as SK466 but plasmid SJK1697 was used. |

| | |
|---|---|
| SK482 | This strain was constructed in two steps. First, we constructed *rne*::*rne-venus* by integrating *venus* into the *rne* region in MG1655 by lambda Red recombination. The *kan* cassette was removed by FLP.<br><br>rne-venus-F:<br>CGGCAACACATCATGCCTCTGCCGCTCCTGCGCGTCCGCAACCTGTTGAGAGA GGTGGTTTATCCAGCAAGG<br>rne-venus-R:<br>AATAAAAAAGCCCTGGCAGTTACCAGGGCTTGATTACTTTGAGCTAATTATCGCT GGTGTAGGCTGGAGC<br><br>Secondly, phage transduction was performed to move *hupA*::*hupA-mCherry-kan* (SK213) into this strain. |
| SK486 | This strain was constructed in two steps, similar to SK482. Only difference is that rne592-Venus-F was used to amplify *venus* when we constructed *rne*::*rne*(1-592)-*venus*. |
| SK505 | Phage transduction of *rne* mutant in SK466 into SK98. |
| SK506 | Phage transduction of *rne* mutant in SK467 into SK98. |
| SK507 | *lacY2-mEos3.2-kan* region was from SK424 using K088 and K140 and inserted into *rne* sequence in MG1655.<br>K140:<br>AATAAAAAAGCCCTGGCAGTTACCAGGGCTTGATTACTTTGAGCTAATTATATCG TGAGGATGCGTCATCG |
| SK508 | Same as SK507 except that the DNA was integrated into SK98 for lambda Red recombination. |
| SK512 | *hupA-mcherry-kan* in CJW5158 was moved to SK290 via phage transduction. |
| SK592 | *lacY6-mEos3.2-kan* region was from SK425 using K088 and K140 and inserted into *rne* sequence in MG1655. |
| SK593 | Same as SK592, but the DNA was integrated into SK98 by lambda Red recombination. |
| SK594 | The *kan* cassette was removed from SK370 by FLP. |
| SK595 | Phage transduction of SK187 into SK98. |
| SK598 | Same as SK466, but plasmid SJK1716 was used for PCR, and the integration occurred into SK594. |
| SK741 | The DNA sequence encoding the mutant MTS (F574AA)-CTD-mEos3.2-Kan was amplified from SK187 using the primers F574AA_for and SJK034. The amplicon was integrated into SK594 by lambda Red recombination.<br>F574AA_for:<br>CTGCACCGGCAACACCAGCAGCTCCTGCACAACCTGGGCTGTTGAGCCGCGCA GCAGGCGCACTGAAAGCGCTGTTCAGC |
| SK742 | The DNA sequence encoding the mutant MTS (F575E)-CTD-mEos3.2-Kan was amplified from SK187 using the primers F575E_for and SJK034. The amplicon was |

| | |
|---|---|
| | integrated into SK594 by lambda Red recombination.<br>F575E_for:<br>CACCGGCAACACCAGCAGCTCCTGCACAACCTGGGCTGTTGAGCCGCTTCGAAGGCGCACTGAAAGCGCTGTTCAGC |
| SK743 | The DNA sequence encoding the mutant MTS (F582E)-CTD-mEos3.2-Kan was amplified from SK187 using the primers F582E_for and SJK034. The amplicon was integrated into SK594 by lambda Red recombination.<br>F582E_for:<br>CTGCACAACCTGGGCTGTTGAGCCGCTTCTTCGGCGCACTGAAAGCGCTGGAAAGCGGTGGTGAAGAAACCAAACC |
| SK748 | The DNA sequence encoding the mutant MTS (F574AA)-mEos3.2-Kan was amplified from the DNA fragment used to construct SK374. It was amplified using the primers F574AA_for and SJK034. The amplicon was integrated into SK98 by lambda Red recombination. |
| SK749 | The DNA sequence encoding the mutant MTS (F575E)-mEos3.2-Kan was amplified from the DNA fragment used to construct SK374. It was amplified using the primers F575E_for and SJK034. The amplicon was integrated into SK98 by lambda Red recombination. |
| SK750 | The DNA sequence encoding the mutant MTS (F582E)-mEos3.2-Kan was amplified from the DNA fragment used to construct SK374. It was amplified using the primers F582E_for and SJK034. The amplicon was integrated into SK98 by lambda Red recombination. |
| **Plasmids** | |
| SJK1606 | We constructed pBAD18kan-Venus-MTS (SJK1591) first by Gibson ligation of 3 DNA fragments. Two fragments were from the plasmid backbone (pBAD18kan[34]), amplified by K042 and aph_in330F and by K061 and aph_in355rev. K061 primer contains the MTS sequence. The third DNA fragment was *venus* sequence amplified from SX701[37] using K046 and K062. K046 contains a proper RBS sequence for translation of *venus*-MTS from the final plasmid. K062 contains a linker sequence between Venus and the MTS.<br>The second Gibson ligation was done with two DNA fragments: linearized plasmid SJK1591 by PCR with K056 and K057 and *mEos3.2* sequence from SK187 by PCR with mEos_5for and mEos_3rev.<br><br>K042: CTAGCCCAAAAAAACGGGTATGG<br>Aph_in330F: CCAGGTATTAGAAGAATATCC<br>K061:<br>CAACCTGGGCTGTTGAGCCGCTTCTTCGGCGCACTGAAAGCGCTGTTCAGCTAACCTGATACAGATTAAATCAGAACG<br>aph_in355rev: CTGAATCAGGATATTCTTCTAATACC<br>K046:<br>CCATACCCGTTTTTTTGGGCTAGTTGGATGGAGTGAAACGATGAGCAAGGGCGAGGAGCTGTTCACC<br>K062:<br>GCCGAAGAAGCGGCTCAACAGCCCAGGTTGGGATAAACCACCTCTTAGCC<br>K056:<br>CACTCGGGCCTGCCGGACAACGCCCGCCGCAAGGGTGGGCGCGCCGACCC<br>K057:<br>GATCTTCATGTCCGGCTTGATCGCCGACATCGTTTCACTCCATCCAACTAGC |

| | |
|---|---|
| | mEos_5for: ATGTCGGCGATCAAGCCGGAC<br>mEos_3rev: GCGGCGGGCGTTGTCCGGCAG |
| SJK1689 | The plasmid was constructed by Gibson ligation of 3 DNA fragments. The first fragment is pUC19 plasmid backbone and *lacI* sequence amplified from plasmid SK141 using two primers: lacZp_rev and lacA_out20F. The second fragment is *lacY2* sequence amplified from MG1655 using primers K117 and K118. The third fragment is CTD-*mEos3.2-kan* sequence amplified from SK187 using rne_in1755F and K124.<br>K117:<br>GATAACAATTTCACACAGGAAACAGCTATGTACTATTTAAAAAACACAAACTTTT GG<br>K118:<br>TGCTGGTTGCTCGGTCGGTTTGGTTTCTTCTTTGCGCAGCCCGAGTTTGTCAGA AAGC<br>rne_in1755F: GAAGAAACCAAACCGACCGAGC<br>K124:<br>GATAAGCGCAGCGTATCAGGCAATTTTTATAATATAAAAAAGCCCTGGCAGTTAC C |
| SJK1697 | The plasmid was constructed in the same way as for SJK1689 except for a different second fragment. It was *lacY6* sequence amplified from MG1655 using primers K117 and K120.<br>K120:<br>TGCTGGTTGCTCGGTCGGTTTGGTTTCTTCAGAAGAGGGCGCATCCGTTTTGG |
| SJK1716 | The plasmid was constructed in the same way as for SJK1689 except for a different second fragment. It was *lacY12* sequence amplified from MG1655 using primers K117 and K123.<br>K123:<br>TGCTGGTTGCTCGGTCGGTTTGGTTTCTTCAGCGACTTCATTCACCTGACG |
| SK567 | The plasmid was constructed by Gibson ligation of two fragments. The first fragment was from pET29b-H6_Streptavidin_sfGFP by digestion with BamHI. The second fragment was *mEos3.2* sequence, amplified from SK187 using primers strep-mEOS-f and strep-mEOS-r.<br>strep-mEOS-f:<br>GAATCCGTTGGACGCTGTCCAACAAGGATC**G**GGATCCGGATCAATGTCGGCGA TCAAGCCGGACATGAAGATCAAGC<br>strep-mEOS-r:<br>GTTCTTCTCCTTTGCTCATTGATCCGGATCCTTAGCGGCGGGCGTTGTCC |

#### Table S3: Doubling times and cell sizes

| Strain Number | Doubling time (min, mean ± std) | Cell length (µm, mean ± std) | Cell width (µm, mean ± std) |
|---|---|---|---|
| SK1 (MG1655) | 82 ± 10 | – | – |
| SK98 | 95.0 ± 9.40 | -- | -- |
| SK187 | 89 ± 9 | 3.362 ± 0.475 | 1.068 ±0.066 |
| SK249 | 106 ± 6 | 3.582 ± 0.547 | 1.053 ±0.067 |

| | | | |
|---|---|---|---|
| SK292 | 83.2 ± 3.2 | 3.287 ± 0.479 | 1.088 ±0.050 |
| SK304 | 93 ± 11 | 3.284 ± 0.516 | 1.083 ±0.054 |
| SK308 | 69.8 ± 1.1 | 3.206 ± 0.479 | 1.083 ±0.050 |
| SK373 | 78 ± 4 | 3.459 ± 0.447 | 1.122 ±0.063 |
| SK374 | 70.8 ± 3.7 | 3.333 ± 0.444 | 1.139 ±0.057 |
| SK404 | 90 ± 7 | 3.27 ± 0.03 | 1.161 ± 0.004 |
| SK411 | 87.6 ± 3.6 | 3.26 ± 0.03 | 1.121 ± 0.004 |
| SK424 | 83 ± 8 | 3.24 ± 0.03 | 1.103 ± 0.004 |
| SK425 | 81 ± 7 | 3.35 ± 0.02 | 1.098 ± 0.003 |
| SK455 | 88 ± 2 | 3.48 ± 0.03 | 1.113 ± 0.002 |
| SK466 | 87 ± 11 | 3.18 ± 0.03 | 1.119 ± 0.003 |
| SK467 | 89 ± 10 | 3.25 ± 0.03 | 1.093 ± 0.004 |
| SK507 | 89 ± 10 | 3.09 ± 0.03 | 1.142 ± 0.005 |
| SK592 | 94.1 ± 0.9 | 3.27 ± 0.02 | 1.188 ± 0.003 |
| SK595 | 98.5 ± 3.3 | -- | -- |
| SK598 | 93.6 ± 7.2 | 3.08 ± 0.02 | 1.050 ± 0.002 |
| SK741 | 94.0 ± 3.6 | 3.49 ± 0.03 | 1.034 ± 0.002 |
| SK742 | 99.7 ± 4.6 | 3.43 ± 0.03 | 1.031 ± 0.003 |
| SK743 | 102.4 ± 3.0 | 3.51 ± 0.03 | 1.047 ± 0.002 |
| SK748 | 94.8 ± 4.0 | 3.59 ± 0.03 | 1.203 ± 0.003 |
| SK749 | 95.2 ± 3.4 | 3.34 ± 0.03 | 1.183 ± 0.004 |
| SK750 | 93.7 ± 3.3 | 3.33 ± 0.02 | 1.161 ± 0.003 |
| SK187<br>(M9 succinate at 30°C) | 153 ± 19 min | 3.37 ± 0.02 | 0.970 ± 0.002 |

**Table S4: xNorm histogram modeling result**

| Protein | Strain number | fcut | | locError (nm) | | dilF | | MB% | |
|---|---|---|---|---|---|---|---|---|---|
| | | mean | std | mean | std | mean | std | mean | stdev |

| | | | | | | | | | |
|---|---|---|---|---|---|---|---|---|---|
| RNE | SK187 | 0.3 | 0.030 | 58.4 | 1.27 | 1.67 | 0.010 | 92.6 | 1.2 |
| LacY | SK292 | 0.25 | 0.006 | 49.3 | 0.52 | 1.62 | 0.002 | 98.5 | 1.0 |
| LacZ | SK407 | 0.36 | 0.020 | 56.3 | 0.92 | 1.87 | 0.020 | 3.4 | 2.1 |
| RNE-F574AA -CTD | SK741 | 0.26 | 0.01 | 55.0 | 0.56 | 1.76 | 0.004 | 88.12 | 1.71 |
| RNE-F582E-CTD | SK743 | 0.28 | 0.01 | 49.1 | 1.05 | 1.9 | 0.002 | 46.84 | 1.43 |
| RNE-F575E-CTD | SK742 | 0.27 | 0.01 | 49.8 | 1.01 | 1.9 | 0.002 | 49.71 | 1.48 |
| RNE ΔMTS | SK249 | 0.29 | 0.01 | 45.4 | 1.26 | 1.9 | 0.003 | 33.71 | 1.33 |
| MTS | SK455 | 0.28 | 0.006 | 47.5 | 0.55 | 1.52 | 0.003 | 99.9 | 0.1 |
| LacY2 | SK424 | 0.26 | 0.005 | 48.3 | 0.68 | 1.58 | 0.003 | 99.6 | 0.4 |
| LacY6 | SK425 | 0.26 | 0.006 | 48.1 | 0.47 | 1.57 | 0.002 | 90.2 | 0.6 |
| RNE ΔCTD or RNE (1-592) | SK374 | 0.27 | 0.004 | 52.4 | 0.52 | 1.56 | 0.002 | 99.8 | 0.2 |
| RNE-LacY2 (ΔCTD) | SK507 | 0.31 | 0.008 | 49.9 | 0.8 | 1.56 | 0.004 | 99.8 | 0.2 |
| RNE-LacY6 (ΔCTD) | SK592 | 0.3 | 0.008 | 45.6 | 0.91 | 1.49 | 0.004 | 99.8 | 0.2 |
| RNE-LacY12 (ΔCTD) | SK404 | 0.27 | 0.006 | 57.1 | 0.6 | 1.55 | 0.002 | 99.3 | 0.6 |
| RNE-LacY2-CTD | SK466 | 0.27 | 0.020 | 55 | 0.64 | 1.7 | 0.004 | 69.4 | 1.9 |
| RNE-LacY6-CTD | SK467 | 0.29 | 0.020 | 56.1 | 0.85 | 1.72 | 0.007 | 86.2 | 1.5 |
| RNE-LacY12-CTD | SK598 | 0.36 | 0.020 | 53.7 | 0.81 | 1.73 | 0.010 | 97.1 | 1.8 |
| RNE in M9succ | SK187 | 0.35 | 0.030 | 59 | 0.87 | 1.86 | 0.020 | 92.1 | 2.7 |
| RNE-LacY2-CTD in M9succ | SK466 | 0.25 | 0.007 | 48 | 1.18 | 1.67 | 0.009 | 43.1 | 1.2 |
| MTS in M9succ | SK455 | 0.32 | 0.040 | 59.8 | 0.92 | 1.67 | 0.020 | 82.8 | 3.4 |
| LacY2 in M9succ | SK424 | 0.29 | 0.006 | 46.7 | 0.49 | 1.7 | 0.004 | 99.9 | 0.1 |
| RNE-F574AA (ΔCTD) | SK748 | 0.31 | 0.0062 | 50.2 | 0.63 | 1.54 | 0.003 | 96 | 0.56 |
| RNE-F582E (ΔCTD) | SK750 | 0.24 | 0.0042 | 54.7 | 0.54 | 1.59 | 0.002 | 67.1 | 0.77 |
| RNE-F575E (ΔCTD) | SK749 | 0.27 | 0.0070 | 52.8 | 0.48 | 1.55 | 0.002 | 91.4 | 0.37 |
| RNE ΔMTS ΔCTD or RNE (1-529) | SK373 | 0.33 | 0.02 | 59.6 | 0.59 | 1.89 | 0.005 | 17.4 | 0.99 |

#### Table S5: qRT PCR primers used in study

These are the same primers as previously described[2].

| Description | Primer sequence (5' to 3') |
|---|---|
| lacZ530F | TTTTACGCGCCGGAGAAAAC |
| lacZ530R | AGTCGGTTTATGCAGCAACG |
| lacZ2732F | TTACTGCCGCCTGTTTTGAC |
| lacZ2732R | TGTAGCGGCTGATGTTGAAC |
| gapA274F | GTTGTCGCTGAAGCAACTGG |
| gapA274R | CGATGTCCTGGCCAGCATAT |

#### Table S6: Figure data statistics

| Figure number | Strain | Strain | Number of tracks/spots | Number of cells |
|---|---|---|---|---|

| | | | | |
|---|---|---|---|---|
| 1D | SK187 | RNE | 143,700 spots | 179 |
| | SK292 | LacY | 199,228 spots | 161 |
| 1F | SK407 | LacZ | 218,120 spots | 247 |
| 2B | SK407 | LacZ | 218,120 spots | 247 |
| | SK249 | RNE ΔMTS | 91,960 spots | 96 |
| | SK373 | RNE ΔMTS ΔCTD | 583,156 spots | 205 |
| 2D | SK187 | WT RNE | 143,700 spots | 179 |
| | SK741 | RNE-F574AA-CTD | 254,252 spots | 417 |
| | SK743 | RNE-F582E-CTD | 284,540 spots | 449 |
| | SK742 | RNE-F575E-CTD | 222,496 spots | 354 |
| | SK249 | RNE ΔMTS | 91,960 spots | 96 |
| 3A | SK187 | RNE, EATA MSD | 11,260 tracks | 177 |
| 3B | SK187 | WT RNE | 11,260 tracks | 177 |
| | SK249 | RNE ΔMTS | 7,539 tracks | 95 |
| | SK374 | RNE ΔCTD | 37,739 tracks | 215 |
| | SK373 | RNE ΔMTS ΔCTD | 36,858 tracks | 205 |
| 3C | SK187 | RNE | 11,260 tracks | 179 |
| | SK187 +rif | RNE +rif | 7,473 tracks | 280 |
| 3D | SK292 | LacY | 20,186 tracks | 159 |
| | SK292 +rif | LacY +rif | 3,364 tracks | 140 |
| 3E | SK47 | L1 | 2,533 tracks | 74 |
| | SK47 +rif | L1 +rif | 1,109 tracks | 51 |
| 4B | SK455 | MTS | 224,352 spots | 577 |
| | SK424 | LacY2 | 107,436 spots | 216 |
| | SK425 | LacY6 | 221,840 spots | 367 |
| | SK292 | LacY12 (full) | 199,228 spots | 161 |
| 4C | SK455 | MTS | 18,804 tracks | 558 |
| | SK424 | LacY2 | 3,812 tracks | 175 |
| | SK425 | LacY6 | 16,009 tracks | 336 |
| | SK292 | LacY12 | 20,186 tracks | 159 |
| 5C | SK292 | LacY | 199,228 spots | 161 |
| | SK374 | RNE ΔCTD | 371,680 spots | 215 |
| | SK507 | RNE-LacY2 ΔCTD | 72,872 spots | 323 |
| | SK592 | RNE-LacY6 ΔCTD | 242,556 spots | 396 |
| | SK404 | RNE-LacY12 ΔCTD | 189,584 spots | 141 |
| 5D | SK292 | LacY | 199,228 spots | 161 |
| | SK187 | WT RNE | 143,700 spots | 179 |
| | SK466 | RNE-LacY2-CTD | 218,416 spots | 250 |
| | SK467 | RNE-LacY6-CTD | 189,672 spots | 180 |
| | SK598 | RNE-LacY12-CTD | 105,656 spots | 450 |
| 5G | SK374 | RNE ΔCTD | 37,739 tracks | 215 |
| | SK507 | RNE-LacY2 ΔCTD | 4,431 tracks | 273 |
| | SK592 | RNE-LacY6 ΔCTD | 11,375 tracks | 371 |
| | SK404 | RNE-LacY12 ΔCTD | 20,263 tracks | 140 |
| 5H | SK187 | WT RNE | 11,260 tracks | 177 |
| | SK466 | RNE-LacY2-CTD | 17,698 tracks | 247 |
| | SK467 | RNE-LacY6-CTD | 11,323 tracks | 176 |
| | SK598 | RNE-LacY12-CTD | 8,060 tracks | 402 |
| S1A | SK187 | RNE | 143,700spots | 179 |
| S1B,C | SK187, live | WT RNE, live | | 179 |
| | SK187, fixed | WT RNE, fixed | | 249 |
| | SK249, live | RNE ΔMTS, live | | 96 |

| | | | | |
|---|---|---|---|---|
| | SK249, fixed | RNE ΔMTS, fixed | | 282 |
| S2A | SK187 | WT RNE | 143,700 spots-all<br>34,040 spots-slow<br>2,376 spots-fast | 179 |
| S2B | SK292 | LacY | 199,228 spots-all<br>60,984 spots-slow<br>888 spots-fast | 161 |
| S2C | SK407 | LacZ | 218,120 spots-all<br>860 spots-slow<br>25,548 spots-fast | 247 |
| S2D | SK249 | RNE ΔMTS | 91,960 spots-all<br>8,720 spots-slow<br>17,816 spots-fast | 96 |
| S2E | SK373 | RNE ΔMTS ΔCTD | 583,156 spots-all<br>21,432 spots-slow<br>104,228 spots-fast | 205 |
| S2F | SK741 | RNE-F574AA-CTD | 254,252 spots-all<br>66,832 spots-slow<br>8,700 spots-fast | 417 |
| S2G | SK743 | RNE-F582E-CTD | 284,540 spots-all<br>23,748 spots-slow<br>42,528 spots-fast | 449 |
| S2H | SK742 | RNE-F575E-CTD | 222,496 spots-all<br>27,588 spots-slow<br>27,420 spots-fast | 354 |
| S2I | SK425 | LacY6 | 221,840 spots-all<br>42,388 spots-slow<br>4,960 spots-fast | 367 |
| S2J | SK466 | RNE-LacY2-CTD | 218,416 spots<br>39,276 spots-slow<br>17,904 spots-fast | 250 |
| S2K | SK467 | RNE-LacY6-CTD | 189,672 spots-all<br>32,508 spots-slow<br>5,100 spots-fast | 180 |
| S2L | SK598 | RNE-LacY12-CTD | 105,656 spots-all<br>24,308 spots-slow<br>696 spots-fast | 450 |
| S2M | SK748 | RNE-F574AA ΔCTD | 541,224 spots-all<br>134,388 spots-slow<br>5,508 spots-fast | 350 |
| S2N | SK750 | RNE-F582E ΔCTD | 489,076 spots-all<br>67,636 spots-slow<br>32,852 spots-fast | 435 |
| S2O | SK749 | RNE-F575E ΔCTD | 259,736 spots-all<br>59,900 spots-slow<br>5,920 spots-fast | 228 |
| S3A | SK187 | RNE | 11,260 tracks | 177 |
| | SK187 +rif | RNE +rif | 7,473 tracks | 280 |
| | SK47 | L1 ribosome | 2,533 tracks | 74 |
| | Protein extracted from plasmid SK567 | His6-streptavidin-mEos3.2 | 10,291 tracks | -- |
| S3B | SK47 | L1 ribosome | 2,533 tracks | 74 |

| | | | | |
|---|---|---|---|---|
| S3C | SK187 | WT RNE | 11,260 tracks | 177 |
| S3F | SK187 | WT RNE | 11,260 tracks | 177 |
| | SK187 +chlor | RNE +chlor | 1,887 tracks | 52 |
| | SK411 | RNE, *lacZ* overexpressed | 15,630 tracks | 279 |
| S8F | SK486, fixed | RNE ΔCTD, fixed | | 133 |
| | SK482, fixed | WT RNE, fixed | | 123 |

#### Table S7: *P*-values determined by two-tailed Student's *t*-test

| Relevant figure | Alternative hypothesis | *p*-value |
|---|---|---|
| 6C | $k_{d1}$ of *lacZ* mRNA in RNE-F574AA-CTD is different from that in WT RNE. | 0.66 |
| | $k_{d1}$ of *lacZ* mRNA in RNE-F575E-CTD is different from that in WT RNE. | 3.8E-04 |
| | $k_{d1}$ of *lacZ* mRNA in RNE-F582E-CTD is different from that in WT RNE. | 0.0076 |
| | $k_{d1}$ of *lacZ* mRNA in RNE-LacY2-CTD is different from that in WT RNE. | 0.021 |
| | $k_{d1}$ of *lacZ* mRNA in RNE-LacY6-CTD is different from that in WT RNE. | 0.60 |
| | $k_{d1}$ of *lacZ* mRNA in RNE-LacY12-CTD is different from that in WT RNE. | 0.41 |
| | $k_{d1}$ of *lacZ* mRNA in RNE ΔMTS is different from that in WT RNE. | 0.013 |
| 6D | $k_{d2}$ of *lacZ* mRNA in RNE-F574AA-CTD is different from that in WT RNE. | 0.12 |
| | $k_{d2}$ of *lacZ* mRNA in RNE-F575E-CTD is different from that in WT RNE. | 0.97 |
| | $k_{d2}$ of *lacZ* mRNA in RNE-F582E-CTD is different from that in WT RNE. | 0.44 |
| | $k_{d2}$ of *lacZ* mRNA in RNE-LacY2-CTD is different from that in WT RNE. | 0.67 |
| | $k_{d2}$ of *lacZ* mRNA in RNE-LacY6-CTD is different from that in WT RNE. | 0.47 |
| | $k_{d2}$ of *lacZ* mRNA in RNE-LacY12-CTD is different from that in WT RNE. | 0.81 |
| | $k_{d2}$ of *lacZ* mRNA in RNE ΔMTS is different from that in WT RNE. | 0.23 |
| 6E | $k_{d1}$ of *lacZ* mRNA in RNE-F574AA ΔCTD is different from that in RNE ΔCTD. | 0.50 |
| | $k_{d1}$ of *lacZ* mRNA in RNE-F575E ΔCTD is different from that in RNE ΔCTD. | 0.041 |
| | $k_{d1}$ of *lacZ* mRNA in RNE-F582E ΔCTD is different from that in RNE ΔCTD. | 0.031 |
| | $k_{d1}$ of *lacZ* mRNA in RNE-LacY2 ΔCTD is different from that in RNE ΔCTD. | 0.58 |
| | $k_{d1}$ of *lacZ* mRNA in RNE-LacY6 ΔCTD is different from that in RNE ΔCTD. | 0.37 |
| | $k_{d1}$ of *lacZ* mRNA in RNE-LacY12 ΔCTD is different from that in RNE ΔCTD. | 0.37 |
| | $k_{d1}$ of *lacZ* mRNA in RNE ΔMTS ΔCTD is different from that in RNE ΔCTD. | 0.034 |
| 6F | $k_{d2}$ of *lacZ* mRNA in RNE-F574AA ΔCTD is different from that in RNE ΔCTD. | 0.18 |
| | $k_{d2}$ of *lacZ* mRNA in RNE-F575E ΔCTD is different from that in RNE ΔCTD. | 0.019 |
| | $k_{d2}$ of *lacZ* mRNA in RNE-F582E ΔCTD is different from that in RNE ΔCTD. | 0.020 |
| | $k_{d2}$ of *lacZ* mRNA in RNE-LacY2 ΔCTD is different from that in RNE ΔCTD. | 0.27 |
| | $k_{d2}$ of *lacZ* mRNA in RNE-LacY6 ΔCTD is different from that in RNE ΔCTD. | 0.14 |

| | | |
|---|---|---|
| | $k_{d2}$ of *lacZ* mRNA in RNE-LacY12 ΔCTD is different from that in RNE ΔCTD. | 0.0037 |
| | $k_{d2}$ of *lacZ* mRNA in RNE ΔMTS ΔCTD is different from that in RNE ΔCTD. | 0.030 |
| S7A | $k_{d1}$ of *lacZ* mRNA in RNase E without mEos3.2 attached and $k_{d1}$ of *lacZ* mRNA in RNase E with mEos3.2 attached are different | 0.9104 |
| | $k_{d2}$ of *lacZ* mRNA when RNase E was without mEos3.2 attached and $k_{d2}$ of *lacZ* mRNA when RNase E has mEos3.2 attached are different | 0.8418 |
| S7B | $k_{d1}$ of *lacZ* mRNA in WT and $k_{d1}$ of *lacZ* mRNA when RNase E is overexpressed are different | 0.9635 |
| | $k_{d2}$ of *lacZ* mRNA in WT is smaller than $k_{d2}$ of *lacZ* mRNA when RNase E is overexpressed | 0.2812 |

#### Table S8: Diffusion coefficient (*D*) and 95% CI

| Strain | Strain | <D> | SEM | <D> from bootstrapping | 95% CI from bootstrapping |
|---|---|---|---|---|---|
| SK47 | Ribosome L1 | 0.053 | 0.002 | 0.0528 | [0.0525, 0.0530] |
| SK47 +rif | Ribosome L1 +rif | 0.37 | 0.008 | 0.372 | [0.371, 0.373] |
| SK187 | WT RNE | 0.018 | 0.0002 | 0.01836 | [0.01834, 0.01838] |
| SK187 +chlor | RNE +chlor | 0.019 | 0.0005 | 0.01902 | [0.01895, 0.01908] |
| SK187 +rif | RNE +rif | 0.027 | 0.0003 | 0.02691 | [0.02687, 0.02694] |
| SK187, M9succ | RNE, M9succ | 0.038 | 0.0002 | 0.03786 | [0.03783, 0.03790] |
| SK249 | RNE ΔMTS | 0.10 | 0.001 | 0.1007 | [0.1005, 0.1008] |
| SK292 | LacY | 0.078 | 0.0004 | 0.07774 | [0.07768, 0.07780] |
| SK292 +rif | LacY +rif | 0.097 | 0.001 | 0.0973 | [0.0972, 0.0975] |
| SK292, M9succ | LacY, M9succ | 0.087 | 0.0003 | 0.08702 | [0.08698, 0.08707] |
| SK373 | RNE ΔMTS ΔCTD | 0.39 | 0.001 | 0.3919 | [0.3917, 0.3921] |
| SK374 | RNE ΔCTD | 0.074 | 0.0003 | 0.07403 | [0.07398, 0.07408] |
| SK374, M9succ | RNE592, M9succ | 0.038 | 0.0002 | 0.03785 | [0.03783, 0.03787] |
| SK404 | RNE-lacY ΔCTD | 0.049 | 0.0004 | 0.04929 | [0.04924, 0.04934] |
| SK407 | LacZ | 0.43 | 0.003 | 0.4322 | [0.4317, 0.4326] |
| SK407, M9succ | LacZ, M9succ | 0.38 | 0.005 | 0.3783 | [0.3777, 0.3790] |
| SK424 | LacY2 | 0.17 | 0.002 | 0.1694 | [0.1691, 0.1698] |
| SK425 | LacY6 | 0.14 | 0.0009 | 0.1367 | [0.1366, 0.1368] |
| SK455 | MTS | 0.11 | 0.0006 | 0.10627 | [0.10618, 0.10636] |
| SK466 | RNE-LacY2-CTD | 0.061 | 0.0007 | 0.06107 | [0.06098, 0.06116] |
| SK467 | RNE-LacY6-CTD | 0.021 | 0.0002 | 0.02094 | [0.02092, 0.02097] |
| SK507 | RNE-LacY2 ΔCTD | 0.098 | 0.002 | 0.0987 | [0.0984, 0.0989] |
| SK592 | RNE-LacY6 ΔCTD | 0.055 | 0.0005 | 0.05463 | [0.05457, 0.05469] |
| SK598 | RNE-LacY12-CTD | 0.017 | 0.0002 | 0.01714 | [0.01712, 0.01717] |
| SK741 | RNE-F574AA-CTD | 0.036 | 0.0003 | 0.03645 | [0.03640, 0.03650] |
| SK742 | RNE-F575E-CTD | 0.094 | 0.0007 | 0.09424 | [0.09415, 0.09433] |
| SK743 | RNE-F582E-CTD | 0.091 | 0.0006 | 0.09135 | [0.09127, 0.09143] |
| SK748 | RNE-F574AA ΔCTD | 0.088 | 0.0004 | 0.08781 | [0.08775, 0.08788] |
| SK749 | RNE-F575E ΔCTD | 0.098 | 0.0007 | 0.09815 | [0.09805, 0.09825] |
| SK750 | RNE-F582E ΔCTD | 0.19 | 0.001 | 0.1944 | [0.1943, 0.1945] |

#### SUPPLEMENTARY FIGURES

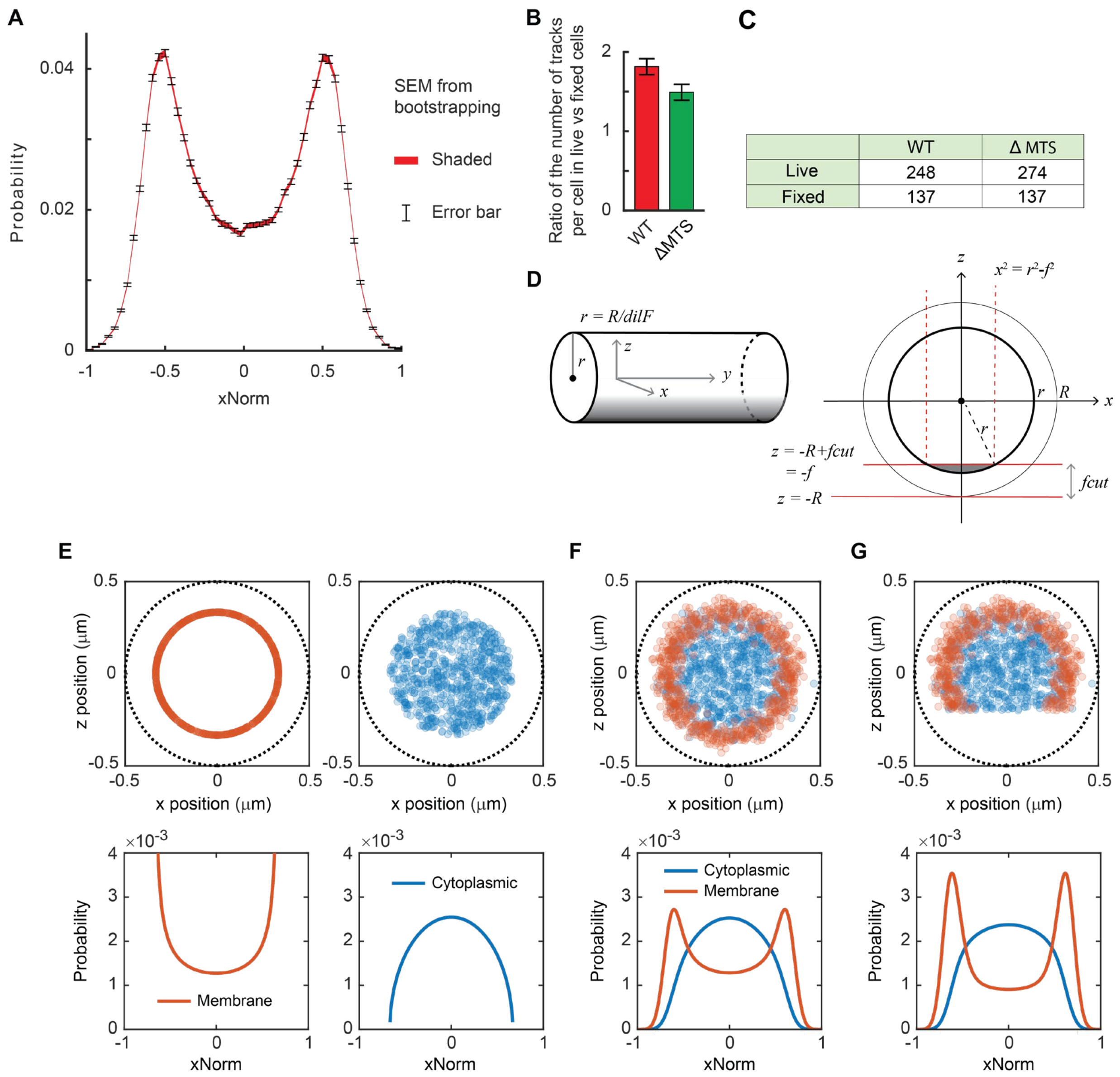

**Figure S1**. Controls for xNorm histograms and MB% calculation. (**A**) The xNorm histogram of RNE shown in **Fig. 1D** is replotted to show SEM in two ways. The SEM from bootstrapping is displayed as error bars (black lines) and the original shaded region (red). This demonstrates that the original presentation with the shaded error region (red) is very small. (**B**) Ratio of the mean number of tracks in live vs fixed cells. WT is for RNE-mEos3.2 (SK187) and ΔMTS is for RNE ΔMTS-mEos3.2 (SK249). Error bars represent the uncertainty derived from the standard deviation of track numbers in live or fixed cells. (**C**) The mean number of tracks per cell measured from live vs fixed cells. Fisher's Exact test result of $p$ = 0.18 suggests no significant association between detectability and cytoplasmic vs membrane localization. (**D**) Coordinates used in a theoretical model for xNorm histogram. (**E**-**G**) Theoretical xNorm histograms for membrane and cytoplasmic molecules. Top row: simulated spot distribution on the membrane (orange) and in the cytoplasm (blue) shown in the vertical cross-section of a cell with diameter of 1 µm. Bottom row: corresponding theoretical xNorm histograms with parameters: (**E**) *dilF* = 1.5, *locErr* = 0, *fCut* = 0, (**F**) *dilF* = 1.5, *locErr* = 40 nm, *fCut* = 0, (**G**) *dilF* = 1.5, *locErr* = 40 nm, *fCut* = 0.3 µm. Random spots were generated either on the periphery of or inside of the circle representing the cell's inner membrane. *dilF* defines the inner membrane away from the cell boundary (black dotted line). In F-G, the localization error was applied by adding a 2D Gaussian noise to all spots. In G, the spots beyond the depth of focus in the z-direction were eliminated. See **Table S6** for data statistics.

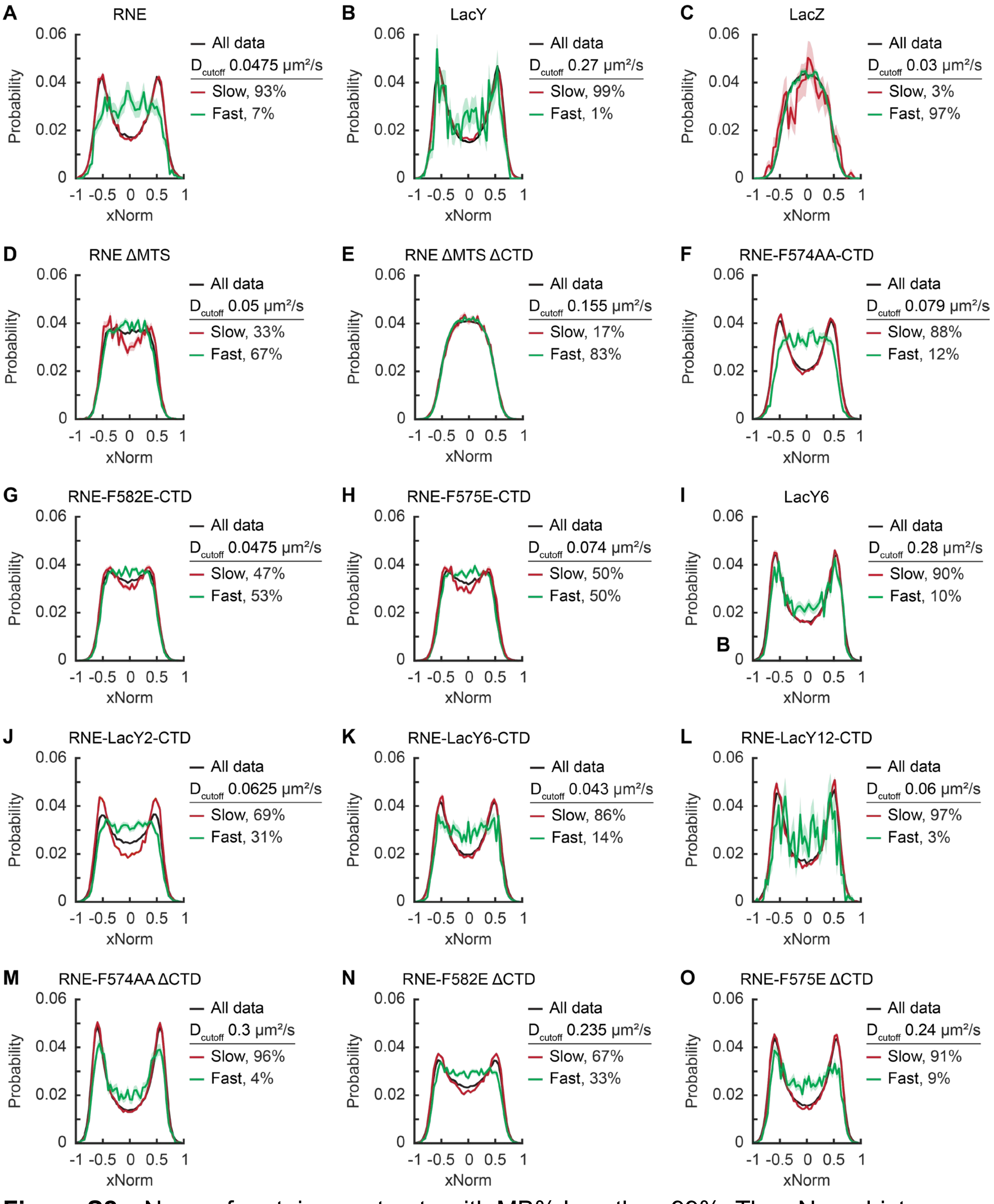

**Figure S2.** xNorm of protein constructs with MB% less than 99%. The xNorm histograms of protein constructs were analyzed by separating molecules (all data) into fast (green) and slow (red) subpopulations. These subpopulations were defined based on the diffusion coefficient *D*: molecules with *D* values in the bottom MB% of the entire population (also below $D_{cutoff}$) were classified as slow, while the other molecules with *D* values above the

$D_{cutoff}$ were classified as fast. Protein constructs with uniform membrane or cytoplasmic localization, such as LacY and LacZ, respectively, showed similar xNorm histograms for both fast and slow populations (e.g., Panels B, C, E, I, L, M). See **Table S6** for data statistics.

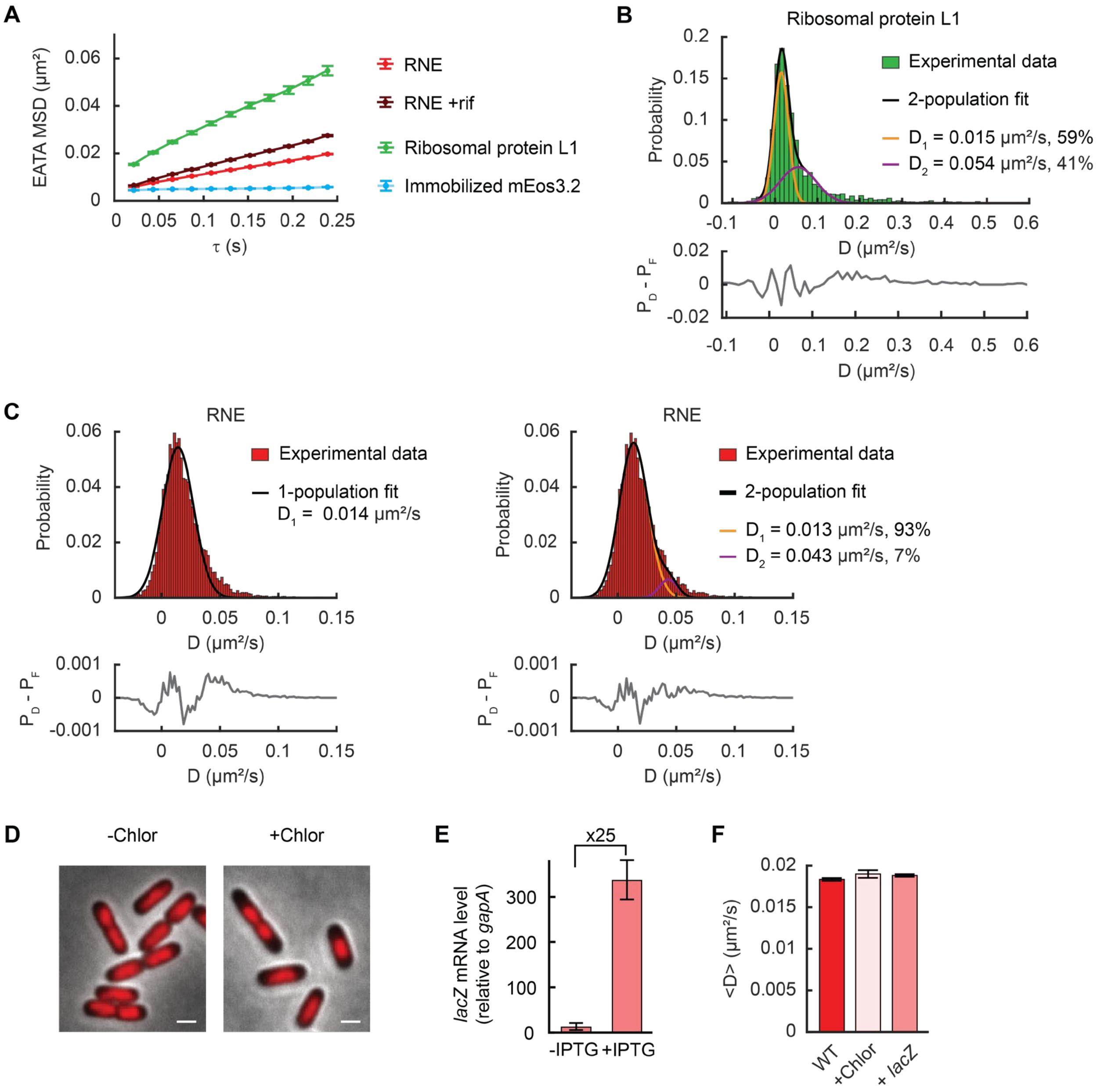

**Figure S3.** Diffusional behavior of RNE and polyribosomes. (**A**) Ensemble-averaged, time-averaged (EATA) MSD of WT RNE, RNE in rif-treated cells, and ribosomal protein L1. Surface-immobilized streptavidin-mEos3.2 was tracked and its EATA MSD was plotted to denote the diffusion of stationary objects. Error bars are the SEM. (**B**) Distribution of ribosomal protein L1's diffusion coefficients and a two-population Gaussian fit ($R^2$ = 0.99). Eq. 4 was used for the fit, yielding $D_1$ of 0.015 [0.014, 0.016] μm$^2$/s and $\sigma_1$ equal to 0.017 [0.016, 0.018] μm$^2$/s for the slow population (polysomes) and $D_2$ equal to 0.054 [0.044, 0.065] μm$^2$/s and $\sigma_2$ of 0.042 [0.037, 0.048] μm$^2$/s for the fast population (free ribosomal units). The fraction of the slow population was 0.59 [0.51, 0.68]. [ ] indicates a 95% confidence interval. (**C**) Distribution of RNE's diffusion coefficients and single- or double-Gaussian fit (both $R^2$ = 0.98). For a single-Gaussian fit, Eq. 3 was used. The fitted *D* coefficient, $D_1$, is 0.014 [0.014, 0.015] μm$^2$/s. The standard deviation of the Gaussian function ($\sigma$) was 0.013 [0.013, 0.014] μm$^2$/s. For a double-Gaussian fit, Eq. 4 was used, yielding $D_1$ of 0.013 [0.013, 0.014] μm$^2$/s and $\sigma_1$ equal to 0.012 [0.012, 0.013] μm$^2$/s for the slow population (membrane-bound) and $D_2$ equal to 0.043 [0.041, 0.0.046] μm$^2$/s and $\sigma_2$ of 0.008 [0.006, 0.01] μm$^2$/s for the fast population (cytoplasmic). The fraction of the slow population was fixed to 93%. (**D**) Nucleoid change upon chloramphenicol treatment. As expected[29], the nucleoid (red, visualized with HU-mcherry, SK512) became more compact (right) compared to untreated cells (left). 100 μg/mL of chloramphenicol was added to the liquid culture for 30 min incubation and also to the agarose pad for imaging. Scale bar = 1 μm. (**E**) Overexpressed *lacZ* mRNA levels measured by qRT-PCR. 1 mM IPTG was added to the exponentially growing SK411 cells for 1 hour. Fold-change was calculated relative to a reference gene, *gapA*. Error bars denote the standard deviation from 3 replicates. (**F**) Mean diffusion coefficient of WT RNE under various cellular conditions. Diffusion of WT RNE was measured in untreated cells (WT; strain SK187), after treating cells with chloramphenicol (100 μg/mL) for 30 min (+Chlor; strain SK187), and in the presence of a high level of *lacZ* mRNA from a high copy plasmid (by treating cells of strain SK411 with 1 mM IPTG for 60 min). Error bars are the SEM. See **Table S6** for data statistics.

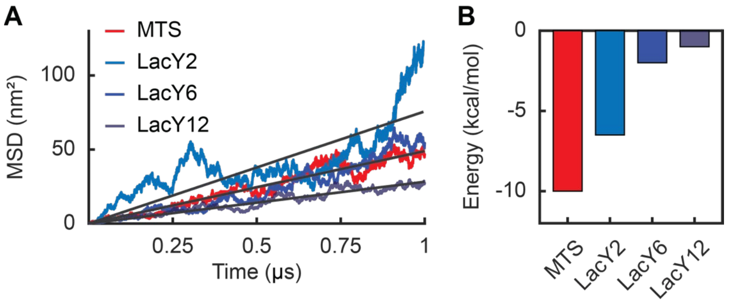

**Figure S4**. All-atom MD simulations of MTS and LacY variants. (**A**) MSD from simulation data. (**B**) Interaction energy between the membrane motif and the lipid membrane.

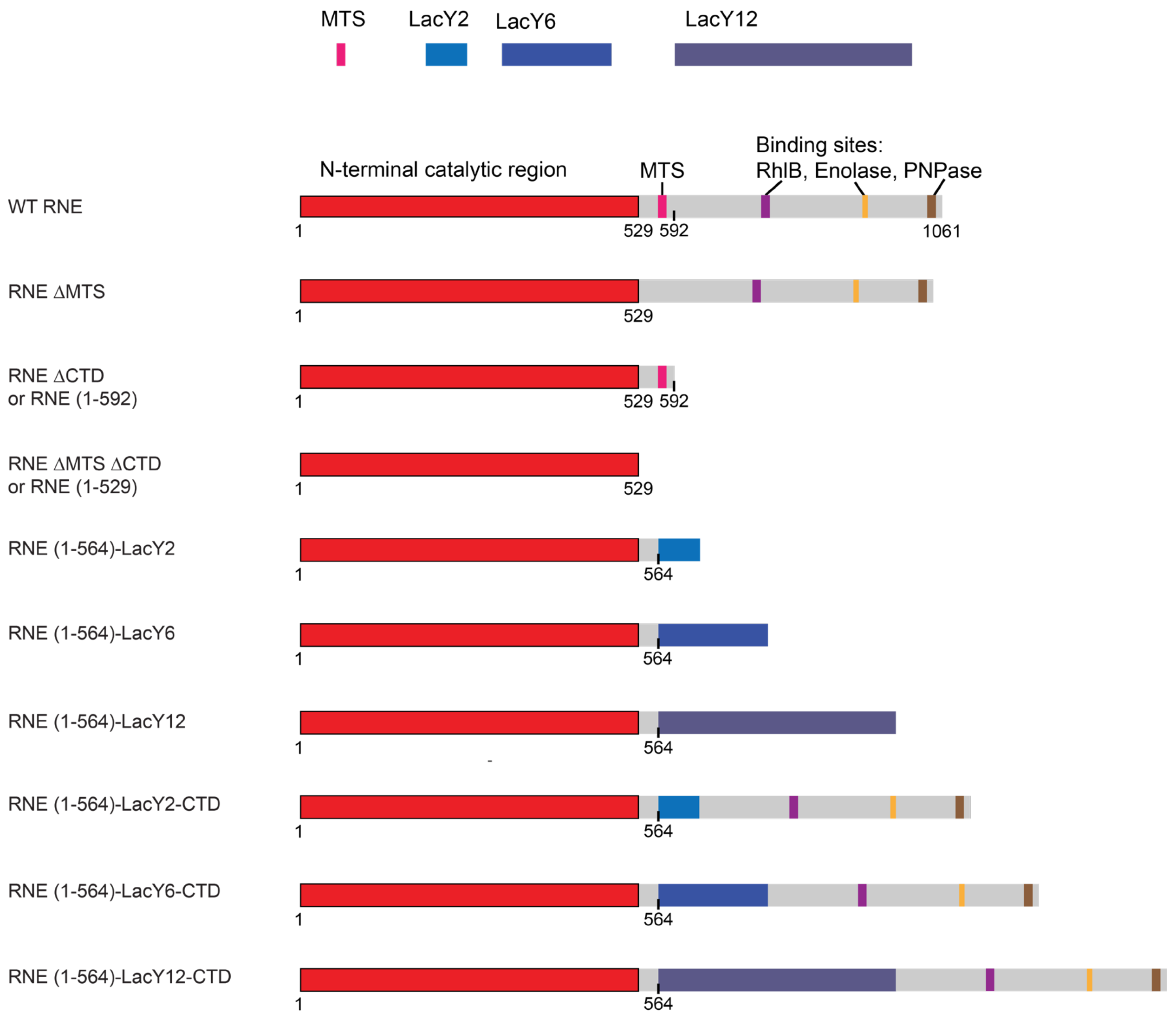

**Figure S5**. Linear representation of RNE monomer in various mutants used in this study. The numbers indicate amino acid residues.

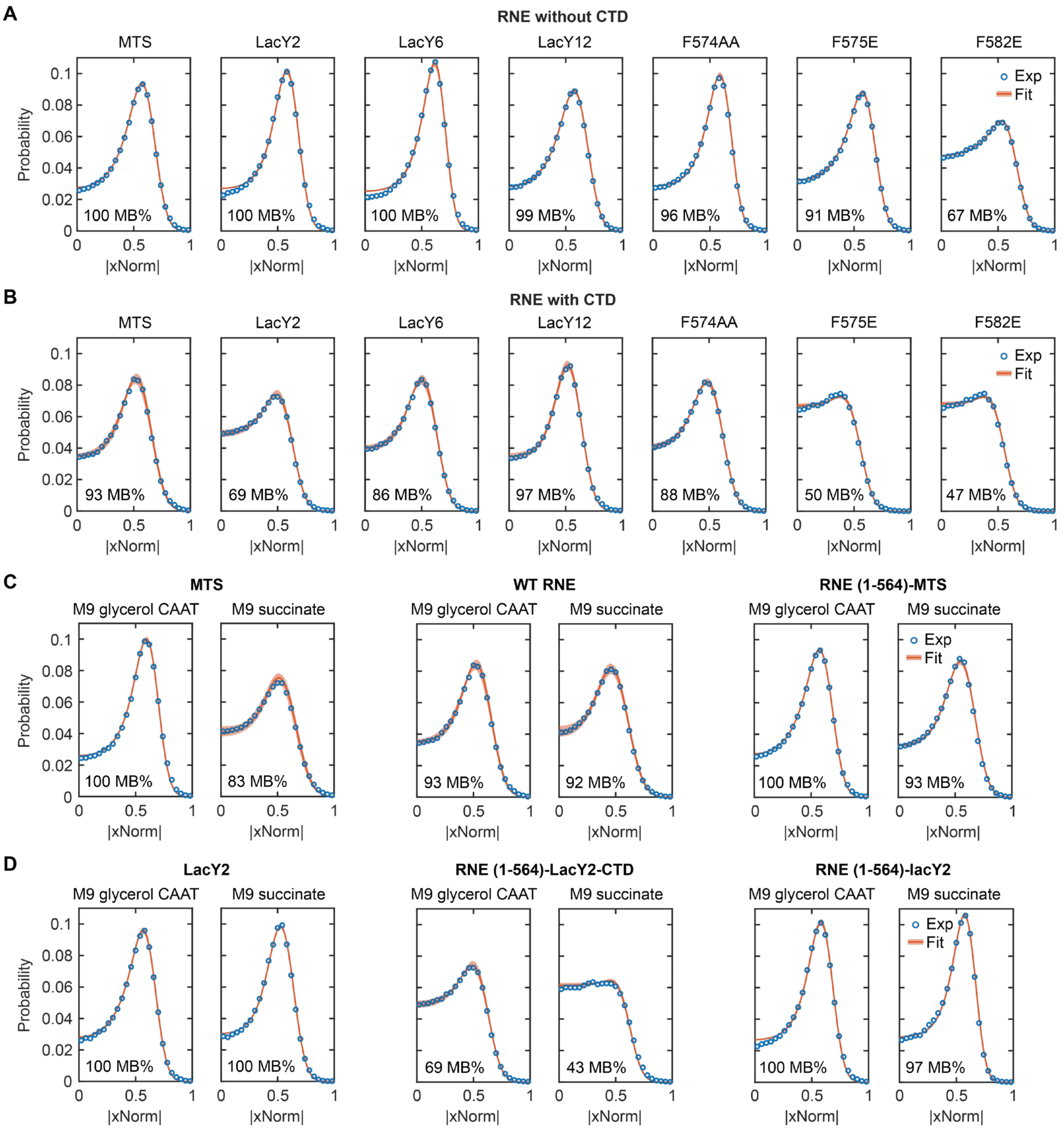

**Figure S6**. xNorm histogram of various RNE mutants used in this study. (**A-B**) Chimeric RNE with various membrane-binding motifs without the CTD (**A**) or with the CTD (**B**). (**C**) Effect of growth media on MB% of the MTS motif, WT RNE, and RNE ΔCTD. (**D**) Effect of growth media on MB% of the LacY2 segment, RNE (1-564)-LacY2-CTD, and RNE (1-564)-LacY2. In all panels, experimental data (blue circles) is compared with MCMC-based fitting result (red). Red shaded regions indicate the expected xNorm histogram based on parameter values within the standard deviation. The MB% shown at the bottom left of each plot is the best-fitting result. See **Table S4** for xNorm histogram modeling results.

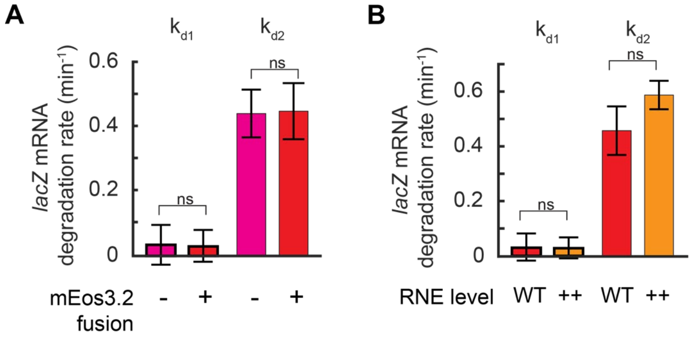

**Figure S7**. *lacZ* mRNA degradation rates under different conditions. (**A**) *lacZ* mRNA degradation rates when RNE is fused to mEos3.2 or not (SK595 vs SK98). (**B**) Degradation rates of nascent mRNA ($k_{d1}$) and released mRNA ($k_{d2}$) of WT RNE (strain SK595) and over-expression of WT RNE (strain SK394). The second-copy RNE was induced with 0.2% arabinose overnight. Error bars are the standard deviations from 3 biological replicates. ns indicates a statistically nonsignificant difference (two-sample *t*-test). See **Table S7** for the *p* values.

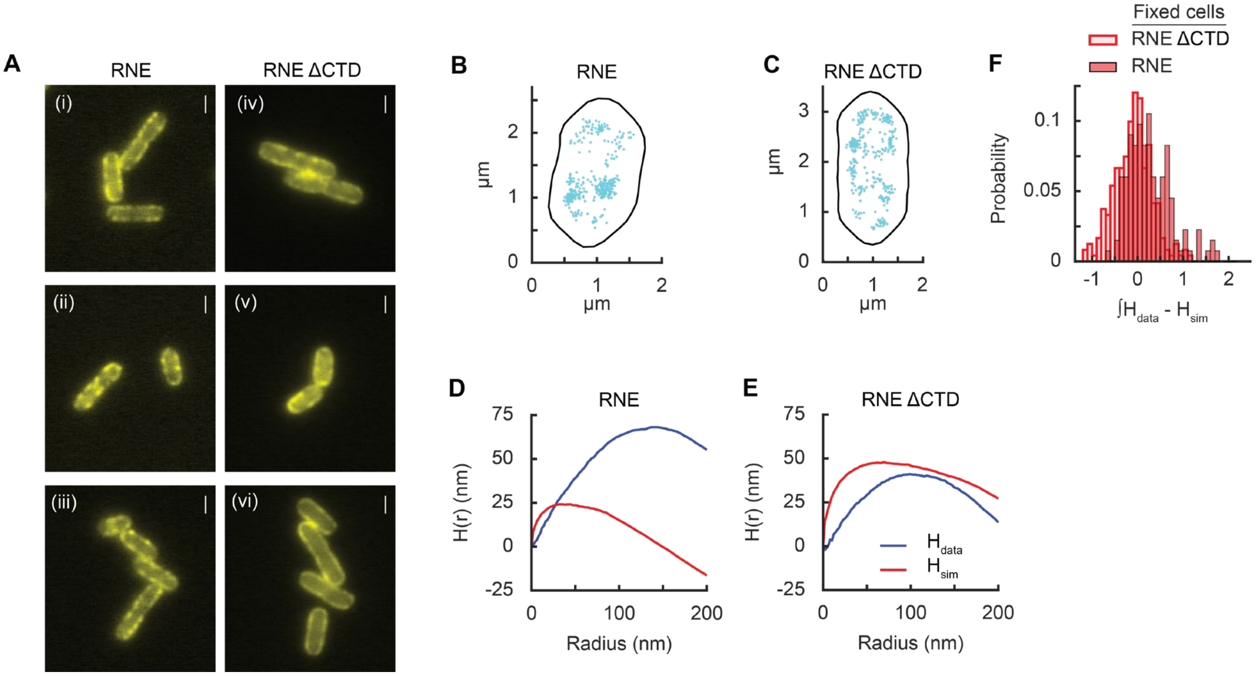

**Figure S8** Clustering of RNE in *E. coli*. (**A**) Example images of WT RNE or RNE ΔCTD (or RNE(1-592)) fused with Venus. The white scale bar is 1 μm. WT RNE (SK482) shows patch localization on the membrane (i-iii). RNE ΔCTD (SK486) also shows patch membrane localization (iv-v), but in some cells (vi), it shows smooth membrane localization. Images are from live cells grown the same way as single-molecule imaging experiments. (**B-C**) RNE-mEos3.2 localization in fixed cells. Each spot represents the first localization event of a track from WT RNE (**B**) or ΔCTD (**C**). (**D**-**E**) H(r) clustering metric for spots shown in panel B and C, respectively. The clustering metric was also calculated for simulated spot distribution based on a complete spatial randomness (CSR) on the membrane with the same number of spots as the experimental cell. (**F**) Cluster analysis based on the difference in the areas of experimental and simulated H(R) data from individual cells (n = 123 cells for WT RNE and n = 133 cells for RNE ΔCTD). RNE ΔCTD is shifted to the left of WT RNE and is statistically different ($p$ = 5.1e-08), indicating RNE ΔCTD is less clustered than WT RNE.

#### REFERENCES

1. Manley, S., Gillette, J. M. & Lippincott-Schwartz, J. in *Methods in Enzymology* Vol. 475 (ed Nils G. Walter) Ch. 5, 109-120 (Academic Press, 2010).
2. Kim, S., Wang, Y.-H., Hassan, A. & Kim, S. Re-defining how mRNA degradation is coordinated with transcription and translation in bacteria. *bioRxiv* (2024). https://doi.org/10.1101/2024.04.18.588412
3. Kim, S. & Vaidya, K. Probing mRNA kinetics in space and time in Escherichia coli using two-color single-molecule fluorescence in situ hybridization. *Journal of Visualized Experiments*, e61520 (2020). https://doi.org/doi:10.3791/61520
4. Paintdakhi, A. *et al.* Oufti: an integrated software package for high-accuracy, high-throughput quantitative microscopy analysis. *Molecular Microbiology* **99**, 767-777 (2016). https://doi.org/https://doi.org/10.1111/mmi.13264
5. Jaqaman, K. *et al.* Robust single-particle tracking in live-cell time-lapse sequences. *Nature Methods* **5**, 695-702 (2008). https://doi.org/10.1038/nmeth.1237
6. Savin, T. & Doyle, P. S. Static and dynamic errors in particle tracking microrheology. *Biophysical Journal* **88**, 623-638 (2005). https://doi.org/10.1529/biophysj.104.042457
7. Saxton, M. J. Single-particle tracking: the distribution of diffusion coefficients. *Biophysical Journal* **72**, 1744-1753 (1997). https://doi.org/https://doi.org/10.1016/S0006-3495(97)78820-9
8. Wieser, S. & Schütz, G. J. Tracking single molecules in the live cell plasma membrane—Do's and Don't's. *Methods* **46**, 131-140 (2008). https://doi.org/https://doi.org/10.1016/j.ymeth.2008.06.010
9. Bakshi, S., Siryaporn, A., Goulian, M. & Weisshaar, J. C. Superresolution imaging of ribosomes and RNA polymerase in live Escherichia coli cells. *Molecular Microbiology* **85**, 21-38 (2012). https://doi.org/https://doi.org/10.1111/j.1365-2958.2012.08081.x
10. Sanamrad, A. *et al.* Single-particle tracking reveals that free ribosomal subunits are not excluded from the Escherichia coli nucleoid. *Proceedings of the National Academy of Sciences* **111**, 11413-11418 (2014). https://doi.org/doi:10.1073/pnas.1411558111
11. Miles, P. R. pymcmcstat: a python package for bayesian inference using delayed rejection adaptive metropolis. *Journal of Open Source Software* **4**, 1417 (2019).
12. Bayas, C. A. *et al.* Spatial organization and dynamics of RNase E and ribosomes in Caulobacter crescentus. *Proceedings of the National Academy of Sciences* **115**, E3712-E3721 (2018). https://doi.org/doi:10.1073/pnas.1721648115
13. Ripley, B. D. Modelling spatial patterns. *Journal of the Royal Statistical Society: Series B (Methodological)* **39**, 172-192 (1977).
14. Besag, J. Discussion on Dr Ripley's Paper. *Journal of the Royal Statistical Society: Series B (Methodological)* **39**, 192-212 (1977).

15 Kiskowski, M. A., Hancock, J. F. & Kenworthy, A. K. On the use of Ripley's K-function and its derivatives to analyze domain size. *Biophysical Journal* **97**, 1095-1103 (2009). https://doi.org/https://doi.org/10.1016/j.bpj.2009.05.039
16 Ehrlich, M. *et al.* Endocytosis by random initiation and stabilization of clathrin-coated pits. *Cell* **118**, 591-605 (2004). https://doi.org/https://doi.org/10.1016/j.cell.2004.08.017
17 Phillips, J. C. *et al.* Scalable molecular dynamics with NAMD. *Journal of Computational Chemistry* **26**, 1781-1802 (2005). https://doi.org/https://doi.org/10.1002/jcc.20289
18 Jo, S., Kim, T., Iyer, V. G. & Im, W. CHARMM-GUI: A web-based graphical user interface for CHARMM. *Journal of Computational Chemistry* **29**, 1859-1865 (2008). https://doi.org/https://doi.org/10.1002/jcc.20945
19 Jorgensen, W. L., Chandrasekhar, J., Madura, J. D., Impey, R. W. & Klein, M. L. Comparison of simple potential functions for simulating liquid water. *The Journal of Chemical Physics* **79**, 926-935 (1983). https://doi.org/10.1063/1.445869
20 Wu, E. L. *et al.* CHARMM-GUI Membrane Builder toward realistic biological membrane simulations. *Journal of Computational Chemistry* **35**, 1997-2004 (2014). https://doi.org/https://doi.org/10.1002/jcc.23702
21 Klauda, J. B. *et al.* Update of the CHARMM all-atom additive force field for lipids: validation on six lipid types. *The Journal of Physical Chemistry B* **114**, 7830-7843 (2010). https://doi.org/10.1021/jp101759q
22 Carpousis, A. J. The RNA degradosome of Escherichia coli: an mRNA-degrading machine assembled on RNase E. *Annual Review of Microbiology* **61**, 71-87 (2007). https://doi.org/https://doi.org/10.1146/annurev.micro.61.080706.093440
23 Mackie, G. A. RNase E: at the interface of bacterial RNA processing and decay. *Nature Reviews Microbiology* **11**, 45-57 (2013). https://doi.org/10.1038/nrmicro2930
24 Blum, E., Py, B., Carpousis, A. J. & Higgins, C. F. Polyphosphate kinase is a component of the Escherichia coli RNA degradosome. *Molecular Microbiology* **26**, 387-398 (1997). https://doi.org/https://doi.org/10.1046/j.1365-2958.1997.5901947.x
25 Carabetta Valerie, J., Silhavy Thomas, J. & Cristea Ileana, M. The response regulator SprE (RssB) is required for maintaining poly(A) polymerase I-degradosome association during stationary phase. *Journal of Bacteriology* **192**, 3713-3721 (2010). https://doi.org/10.1128/jb.00300-10
26 Jaso-Vera, M. E., Domínguez-Malfavón, L., Curiel-Quesada, E. & García-Mena, J. Dynamics of the canonical RNA degradosome components during glucose stress. *Biochimie* **187**, 67-74 (2021). https://doi.org/https://doi.org/10.1016/j.biochi.2021.05.006
27 Callaghan, A. J. *et al.* Structure of Escherichia coli RNase E catalytic domain and implications for RNA turnover. *Nature* **437**, 1187-1191 (2005). https://doi.org/10.1038/nature04084
28 Spring, T. G. & Wold, F. The purification and characterization of Escherichia coli enolase. *Journal of Biological Chemistry* **246**, 6797-6802 (1971). https://doi.org/https://doi.org/10.1016/S0021-9258(19)45916-4
29 Stracy, M. *et al.* Live-cell superresolution microscopy reveals the organization of RNA polymerase in the bacterial nucleoid. *Proceedings of the National Academy of Sciences* **112**, E4390-E4399 (2015). https://doi.org/doi:10.1073/pnas.1507592112
30 Reyer, M. A. *et al.* Kinetic modeling reveals additional regulation at co-transcriptional level by post-transcriptional sRNA regulators. *Cell Reports* **36**, 109764 (2021). https://doi.org/https://doi.org/10.1016/j.celrep.2021.109764
31 Strahl, H. *et al.* Membrane recognition and dynamics of the RNA degradosome. *PLOS Genetics* **11**, e1004961 (2015). https://doi.org/10.1371/journal.pgen.1004961
32 Kim, S., Beltran, B., Irnov, I. & Jacobs-Wagner, C. Long-distance cooperative and antagonistic RNA polymerase dynamics via DNA supercoiling. *Cell* **179**, 106-119.e116 (2019). https://doi.org/https://doi.org/10.1016/j.cell.2019.08.033

33 Thappeta, Y. *et al.* Glycogen phase separation drives macromolecular rearrangement and asymmetric division in E. coli. *bioRxiv* (2024). https://doi.org/10.1101/2024.04.19.590186

34 Guzman, L. M., Belin, D., Carson, M. J. & Beckwith, J. Tight regulation, modulation, and high-level expression by vectors containing the arabinose PBAD promoter. *Journal of Bacteriology* **177**, 4121-4130 (1995). https://doi.org/doi:10.1128/jb.177.14.4121-4130.1995

35 Datsenko, K. A. & Wanner, B. L. One-step inactivation of chromosomal genes in Escherichia coli K-12 using PCR products. *Proceedings of the National Academy of Sciences* **97**, 6640-6645 (2000). https://doi.org/doi:10.1073/pnas.120163297

36 Khemici, V., Poljak, L., Luisi, B. F. & Carpousis, A. J. The RNase E of Escherichia coli is a membrane-binding protein. *Molecular Microbiology* **70**, 799-813 (2008). https://doi.org/10.1111/j.1365-2958.2008.06454.x

37 Choi, P. J., Cai, L., Frieda, K. & Xie, X. S. A stochastic single-molecule event triggers phenotype switching of a bacterial cell. *Science* **322**, 442-446 (2008). https://doi.org/doi:10.1126/science.1161427
